# Characterization of the mycobiome of the seagrass, *Zostera marina*, reveals putative associations with marine chytrids

**DOI:** 10.1101/735050

**Authors:** Cassandra L. Ettinger, Jonathan A. Eisen

**Affiliations:** Genome Center, University of California, Davis, CA, United States; Department of Evolution and Ecology, University of California, Davis, CA, United States; Department of Medical Microbiology and Immunology, University of California, Davis, Davis, CA, United States

**Keywords:** seagrass, *Zostera marina*, mycobiome, marine fungi, ITS, Lobulomycetales, Chytridiomycota

## Abstract

Seagrasses are globally distributed marine flowering plants that are foundation species in coastal ecosystems. Seagrass beds play essential roles as habitats and hatcheries, in nutrient cycling and in protecting the coastline from erosion. Although many studies have focused on seagrass ecology, only a limited number have investigated their associated fungi. In terrestrial systems, fungi can have beneficial and detrimental effects on plant fitness. However, not much is known about marine fungi and even less is known about seagrass associated fungi. Here we used culture-independent sequencing of the ribosomal internal transcribed spacer (ITS) region to characterize the taxonomic diversity of fungi associated with the seagrass, *Zostera marina*. We sampled from two *Z. marina* beds in Bodega Bay over three time points to investigate fungal diversity within and between plants. Our results indicate that there are many fungal taxa for which a taxonomic assignment cannot be made living on and inside *Z. marina* leaves, roots and rhizomes and that these plant tissues harbor distinct fungal communities. The most prevalent ITS amplicon sequence variant (ASV) associated with *Z. marina* leaves was classified as fungal, but could not initially be assigned to a fungal phylum. We then used PCR with a primer targeting unique regions of the ITS2 region of this ASV and an existing primer for the fungal 28S rRNA gene to amplify part of the 28S rRNA gene region and link it to this ASV. Sequencing and phylogenetic analysis of the resulting partial 28S rRNA gene revealed that the organism that this ASV comes from is a member of Novel Clade SW-I in the order Lobulomycetales in the phylum Chytridiomycota. This clade includes known parasites of freshwater diatoms and algae and it is possible this chytrid is directly infecting *Z. marina* leaf tissues. This work highlights a need for further studies focusing on marine fungi and the potential importance of these understudied communities to the larger seagrass ecosystem.

## Introduction

Seagrasses are fully submerged marine flowering plants (angiosperms) that play essential roles in marine ecosystems as foundation species. Although angiosperms are the most diverse terrestrial plant group with over 250,000 species, there are only around 70 species of seagrasses. There are three main lineages of seagrass in the order Alismatales that separately moved into and adapted to the marine ecosystem through convergent evolution between 70 and 100 million years ago (Les et al., 1997; Wissler et al., 2011). Seagrasses are important keystone species in most coastal environments around the world with ecosystem services comparable to those of tropical rainforests (Costanza et al., 1997). However, seagrass beds are increasingly impacted by climate change, pollution and habitat fragmentation and restoration is expensive and has a low success rate (Orth et al., 2006). Although many studies have focused on the ecological importance of seagrasses, relatively little is known about the fungi associated with these species.

Fungi are known to affect land plant fitness both in detrimental ways (e.g., as pathogens) and beneficial ways (e.g., mycorrhizae are involved in facilitating phosphorus and nitrogen uptake for their plant hosts (Bonfante and Anca, 2009)). It is estimated that 82-85% of angiosperm species have mycorrhizal fungal associations (Radhika and Rodrigues, 2007; Wang and Qiu, 2006). Mycorrhizae were previously thought to not colonize aquatic environments, but have since been found in wetlands, estuaries, mangrove forests and freshwater ecosystems (Beck-Nielsen and Madsen, 2001; Bohrer et al., 2004; Kohout et al., 2012; Radhika and Rodrigues, 2007; Šraj-Kržič et al., 2006). Mycorrhizal associations are believed to be at least 400 million years old, critical for plant terrestrialization (Humphreys et al., 2010; Redecker et al., 2000; Wang et al., 2010) and the ancestral state for angiosperms (Brundrett, 2002).Despite their importance to land plants, seagrasses are believed to not form mycorrhizal associations (Nielsen et al., 1999), even though freshwater relatives of seagrasses have been observed to sporadically form mycorrhizal associations (Beck-Nielsen and Madsen, 2001; Brundrett, 2009; Clayton and Bagyaraj, 1984; Khan and Belik, 1995; Radhika and Rodrigues, 2007; Šraj-Kržič et al., 2006; Wang and Qiu, 2006; Wigand and Court Stevenson, 1994).

In contrast to their apparent lack of mycorrhizal associations, some studies of seagrasses have observed associations with novel fungal endophytes similar to dark septate endophytes (DSE) common in land plants (Borovec and Vohník, 2018; Torta et al., 2015; Vohník et al., 2015, 2016, 2017, 2019). DSE are a morphology based type and not a phylogenetic group, and are largely uncharacterized. In some cases, DSEs have been shown to transfer nitrogen and receive carbon from plants as well as increase overall plant nutrient content and growth (Porras-Alfaro and Bayman, 2011; Usuki and Narisawa, 2007). DSEs are not the only fungi that have been observed to form associations with seagrasses. Additional culture based studies have found fungi associated with the leaves, roots and rhizomes of different seagrasses; however, many of these studies often conflict on the prevalence and taxonomic identities of the fungi observed and suffer from the limitations of culture-based methods (Cuomo et al., 1985; Devarajan and Suryanarayanan, 2002; Gnavi et al., 2014; Kirichuk and Pivkin, 2015; Kuo, 1984; Ling et al., 2015; Mata and Cebrián, 2013; Newell, 1981; Panno et al., 2013; Phongpaichit and Supaphon, 2014; Sakayaroj et al., 2010; Shoemaker and Wyllie-Echeverria, 2013; Supaphon et al., 2013, 2017; Torta et al., 2015; Venkatachalam, 2015; Venkatachalam et al., 2015; Vohník et al., 2016). This is compounded by the fact that marine fungi are understudied and that marine fungal culture collections are limited with only around ∼1100 cultured isolates, despite estimates that there are 10,000 or more marine fungal species (Amend et al., 2019; Jones et al., 2015; Jones and Gareth Jones, 2011). Here, we use culture-independent methods to characterize the taxonomic diversity of fungi associated with the seagrass, *Z. marina*, in order to build a framework from which we can begin to determine the evolutionary, ecological, and functional importance of these associations.

## Methods

### Sample collection

Roots, leaves and rhizome tissues from individual *Z. marina* plants and adjacent sediment were collected by coring into the sediment using a 2.375 inch diameter modified PVC pipe (McMaster-Carr, Elmhurst, IL, USA). PVC pipes were modified such that one end of the pipe was cut at an angle to make insertion into the sediment easier. Samples were collected from two sites in Bodega Bay, CA, Gaffney Point (GPS: 38°18’45.05”N, 123° 3’16.66”W) and Westside Point (GPS: 38°19’10.67”N, 123° 3’13.71”W), at three timepoints two weeks apart spanning July-August 2016. Samples from GP were collected only at the first timepoint. Samples were dissected using sterile scissors into different bulk sample types (e.g. roots, leaves, rhizome tissues) in the field, placed into 1.5 mL centrifuge tubes which were immediately placed on dry ice for transport and then kept at −80 °C until analyzed. Sediment was collected from the 2.375 inch PVC pipe using an 11 mm diameter straw which was placed directly next to the individual seagrass shoot tissue in order to assure sediment was collected as close to the plant as possible. For the third timepoint (T3), multiple samples were collected from the same plant. Leaf tissue was cut into ∼5 inch long segments moving along the length of the leaf tissue, individual roots were collected and an additional 7 mm diameter straw core was obtained and then dissected into ∼1.5 cm sections of sediment. Cores of unvegetated sediment from between sites were obtained as a biological control at each time point. For sample sizes, refer to Tables S1-3.

### Molecular methods

DNA was extracted from samples with the PowerSoil DNA Isolation kit (MO BIO Laboratories, Inc., Carlsbad, CA, USA) with minor changes to the manufacturer’s protocol as follows. To improve fungal lysis, samples were heated at 70 °C for 10 minutes between steps 4 and 5 as suggested by the manufacturer. For step 5, samples were bead beaten on the homogenize setting for 1 minute using a BioSpec Products mini-bead beater. Samples were placed into randomized blocks prior to extraction using a random number generator.

In order to obtain loosely associated, or epiphytic, fungi, tissue samples were washed in 1 mL of a 1:50 dilution of Redford buffer solution (1 M Tris-HCL, 0.5 M NaEDTA, 1.2% Triton-X) (Kadivar and Stapleton, 2003; Kembel et al., 2014). Samples were left in this solution for 5 minutes during which they were periodically vortexed. Tissues were then removed and placed into a new microcentrifuge tube for use in endophyte analyses. The epiphyte wash solution was centrifuged at 4000 g for 20 minutes and the pellets were resuspended in MoBio PowerSoil C1 solution, the first step in the DNA extraction process.

After being washed to obtain epiphytes, a subset of samples collected at the third sampling timepoint were further surface cleaned with the goal of removing any leftover loosely associated fungi. This was done in an effort to obtain samples containing only endophytes, which we define here as being tightly associated to the outside or inside the tissue. Tissues were rinsed with autoclaved nanopure water (30 seconds), immersed in 95% ethanol (5 seconds), immersed in 0.5% NaOCl (∼10% bleach) (2 minutes), immersed in 70% ethanol (2 minutes), then rinsed with autoclaved nanopure water (1 minute) three times. Tissues were then placed into MoBio PowerSoil C1 solution and crushed using flame sterilized tweezers.

### Sequence generation

Using a random number generator, samples were randomly assigned places in 96-well plates and sent to Zymo Research, Inc for sequencing via their ZymoBIOMICS Service. We briefly summarize their protocol here. Fungal ITS2 amplicon sequencing was performed using the Quick NGS Library Preparation Kit (Zymo Research, Irvine, CA). The ITS2 region was amplified via PCR using the ITS5.8S_Fun and ITS4_Fun (McHugh and Schwartz, 2015; Taylor et al., 2016) primers which were chosen to try to minimize the amount of host plant amplification. Final PCR products were quantified with qPCR fluorescence readings and pooled together based on equal molarity. The final pooled library was cleaned up with Select-a-Size DNA Clean & Concentrator™ (Zymo Research, Irvine, CA), then quantified with TapeStation (Agilent, Santa Clara, CA) and Qubit (Invitrogen, Carlsbad, CA). The final library was sequenced on Illumina MiSeq (Illumina, Inc., San Diego, CA, USA) with a v3 reagent kit (600 cycles) to generate 300 bp paired-end reads. The sequencing was performed with >10% PhiX spike-in. Sequence reads were demultiplexed by Zymo Research, Inc using the MiSeq Reporter. The raw sequence reads generated for this ITS amplicon project were deposited at GenBank under accession no. <u>PRJNA515720</u>.

### Sequence processing

Primers were removed using cudadapt (v. 1.14) (Martin, 2011). Forward and reverse reads were then merged with PEAR (v. 0.9.5) (Zhang et al., 2014) using the script run_pear.pl (Comeau et al., 2017) which implements PEAR in parallel.

The resulting merged fastq files were analyzed in R (v. 3.5.1) using dada2 (v. 1.10.0), decontam (v. 1.1.2), phyloseq (v. 1.26.0), vegan (v. 2.5.5), FSA (v. 0.8.25) and ggplot2 (v. 3.1.0) (Callahan et al., 2016; Davis et al., 2018; Dixon, 2003; McMurdie and Holmes, 2013; Ogle, 2016; R Core Team, 2016; Wickham, 2009). For a detailed walkthrough of the following analysis using R, see the R-markdown summary file (File S1).

Prior to denoising, reads were truncated at the first quality score of 2 and reads with an expected error greater than 2 were removed. Since PEAR alters the quality scores during sequence merging, we inflated the error profile used by dada2 to denoise sequences by a factor of 3. Reads were then denoised and merged using dada2 to generate a table of amplicon sequence variants (ASVs). Chimeric sequences were identified using removeBimeraDenovo and removed prior to downstream analyses (∼0.31% of sequences). Taxonomy was inferred using the RDP Naive Bayesian Classifier algorithm with a modified UNITE (v. 8.0) database resulting in 2953 ASVs (UNITE Community, 2019; Wang et al., 2007). The UNITE database was modified to include a representative ITS amplicon sequence for the host plant, *Zostera marina* (KM051458.1). ASVs were then named by giving each a unique number preceded by “SV” which stands for sequence variant (e.g. SV1, SV2, etc).

Subsequently, ITS-x (v. 1.1b) was run on the unique ASVs (Bengtsson-Palme et al., 2013). This program, which uses kingdom-specific hidden markov models to identify the ITS region of sequences, could not detect a fungal ITS region in 1064 of the ASVs. These ASVs were thus considered putively non-fungal and were removed from downstream analysis.

Decontam’s prevalence method was used to identify possible contaminants with a threshold of 0.5 which will identify sequences that have a higher prevalence in negative controls than in true samples. This threshold identified 38 possible contaminants which were then removed from the dataset. Negative and positive controls were subsequently removed at this point in the analysis. Remaining ASVs assigned as non-fungal at the domain level (e.g. ASVs assigned to the host plant, *Z. marina*, or with no domain level classification) were removed from the dataset prior to downstream analysis resulting in a final table of 1850 ASVs. Four samples were represented by zero sequences after these filtering steps and were subsequently removed.

### Sequence analysis and visualization

For statistical analysis and visualization, the resulting data was then subset without replacement to an even number of sequences per sample depending on the comparisons being made. As a result, when investigating differences in epiphytes between bulk sample types a depth of 10,000 sequences was selected resulting in the inclusion of 49 samples: leaf (n = 13), root (n = 14), rhizome (n = 7) and sediment (n = 15). When investigating differences across leaf length a number of 5,000 sequences was chosen resulting in the inclusion of 50 samples: leaf epiphytes (n = 25) and leaf endophytes (n = 25). These depths were selected after examining rarefaction curves to balance maximizing the number of sequences per sample while also minimizing the number of samples removed from downstream analysis.

To assess within-sample (i.e., alpha) diversity, the observed number of ASVs and the Shannon index of samples were calculated using the estimate_richness function in phyloseq. Kruskal– Wallis tests with 9,999 permutations were used to test for significant differences in alpha diversity between bulk sample types (leaf, root, rhizome, sediment). For metrics in which the Kruskal–Wallis test resulted in a rejected null hypothesis (*p* < 0.05), Bonferroni corrected post-hoc Dunn tests were performed.

To assess between-sample (i.e., beta), diversity, Bray–Curtis (Bray et al., 1957) dissimilarities were calculated using the ordinate function in phyloseq and visualized using principal coordinates analysis. To test for significant differences in mean centroids between sample categories (e.g., sample type), permutational manovas (PERMANOVAs) were performed with 9,999 permutations and to account for multiple comparisons, p-values were adjusted using the Bonferroni correction (Anderson, 2001). PERMANOVA tests are known to be sensitive to differences in dispersion when using abundance-based distance matrices like Bray-Curtis (Warton et al., 2012), but are still more robust than other tests (Anderson and Walsh, 2013). To control for this, we also tested for differences in mean dispersions between different sample categories using the betadisper and permutest functions from the vegan package in R with 9,999 permutations. For betadisper results that resulted in a rejected null hypothesis (*p* < 0.05), the post-hoc Tukey’s Honest Significant Difference (HSD) test was performed to identify which categories had mean dispersions that were significantly different.

To compare fungal community composition, we collapsed ASVs into taxonomic orders using the tax_glom function in phyloseq and then removed orders with a variance of less than one percent when comparing between sample types and a variance of less than 0.1 percent when comparing across leaf lengths. The average relative abundance of taxonomic orders was compared between sample types using Bonferroni corrected Kruskal–Wallis tests in R. For orders where the Kruskal–Wallis test resulted in a rejected null hypothesis, Bonferroni corrected post-hoc Dunn tests were performed to identify which sample comparisons for each taxonomic order were significantly different.

To examine the contribution of specific ASVs to fungal community composition across leaf length, the dataset was filtered to include only ASVs with a mean abundance of greater than two percent. The resulting ASVs were then compared to each other and those that shared greater than 99% sequence identity were grouped into what we refer to as “complexes”. Complexes were given a name based on the the most abundant ASV in the group (e.g. SV8 complex). The average relative abundance of ASVs and ASV complexes were compared between sample types using Bonferroni corrected Kruskal–Wallis tests in R. For ASVs and ASV complexes where the Kruskal–Wallis test resulted in a rejected null hypothesis, Bonferroni corrected post-hoc Dunn tests were performed to identify which sample comparisons for each ASV or ASV complex were significantly different.

### Sanger sequencing

The most prevalent sequences associated with *Z. marina* leaf tissue all come from a single complex (i.e., they all share > 99% sequence identity). This complex, which includes SV8, SV11, SV16 and SV56, has been named the SV8 complex because SV8 is the most abundant member of this group. None of the ASVs in the SV8 complex could be classified into a fungal phylum using currently available databases. In order to attempt to improve taxonomic classification of the SV8 complex, we used PCR to obtain sequence data for part of the 28S rRNA gene from representatives of this complex found in our samples. To obtain this sequence, a SV8-specific primer (5’-GGAGCATGTCTGTTTGAGAA-3’) was designed for the ITS2 region using Primer-BLAST (Ye et al., 2012). This primer was then used in PCR along with the reverse fungal 28S rRNA gene primer, LR3 (Vilgalys and Hester, 1990). PCR using this pair of primers should at least in theory amplify the ITS2 and D1/D2 regions of the 28S rRNA gene for the SV8 complex.

Using these primers, we performed PCR on DNA from two leaf epiphyte samples (sample IDs: 108A and 109A) using Taq DNA Polymerase (QIAGEN, Hilden, Germany) with the following conditions: 95°C for 5 minutes, 35 cycles at 94°C for 30 seconds, 52°C for 15 seconds, 72°C for 1 minute, and a final extension at 72°C for 8 minutes (Kress and Erickson, 2012).

PCR products were purified using the Nucleospin Gel and PCR kit (QIAGEN, Hilden, Germany). The resulting amplicon was sequenced using the Sanger method by the College of Biological Sciences ^UC^DNA Sequencing Facility (http://dnaseq.ucdavis.edu/). The resulting ABI files were viewed and a consensus sequence was produced using seqtrace following the Swabs to Genomes workflow (Dunitz et al., 2015). Consensus sequences for the PCR products were deposited to NCBI Genbank under accession no. <u>MK994004, MK994005</u>.

### Phylogenetic reconstruction

Closely related sequences to the sequence from the PCR products above were identified using NCBI’s Standard Nucleotide BLAST’s megablast option with default settings. These results were then used to guide a literature search to identify additional sequences for inclusion during phylogenetic reconstruction (Table S4) (Hassett et al., 2017; Karpov et al., 2014; Rad-Menéndez et al., 2018; Simmons et al., 2009; Van den Wyngaert et al., 2018).

A sequence alignment of all sequences listed in Table S4 was generated using MAFFT (v. 7.402) (Katoh, 2002) with default parameters on the CIPRES Science Gateway web server (Miller et al., 2010). The alignment was trimmed using trimAl (v.1.2) with the-gappyout method (Capella-Gutiérrez et al., 2009). The resulting alignment included 22 sequences and 915 alignment positions. JModelTest2 (v. 2.1.6) was run with default parameters on the CIPRES Science Gateway web server to select a best-fit model of nucleotide substitution for use in phylogenetic analyses (Darriba et al., 2012; Guindon and Gascuel, 2003). The best-fit model based on both the Akaike Information Criterion and Bayesian Information Criterion values was the GTR + G + I evolutionary model.

Using the CIPRES Science Gateway web server, phylogenetic trees were inferred from the trimmed alignment using both Bayesian and maximum likelihood approaches and a GTR + G + I model. Bayesian phylogenetic inference was performed using MrBayes (v. 3.2.2) with four incrementally heated simultaneous Monte Carlo Markov Chains (MCMC) run over 40,000 generations, which was the optimal number of generations required to achieve a stop value of 0.01 or less for the convergence diagnostic (Huelsenbeck and Ronquist, 2001). The first 25% of trees generated were discarded as burn-in and for the remaining trees, a majority rule consensus tree was generated and used to calculate the Bayesian Posterior Probabilities. A maximum likelihood approach was undertaken using RAxML (v. 8) using the-autoMRE option which automatically determines the optimal number of bootstrap replicates (Stamatakis, 2014). The resulting phylogenies were then visualized with FigTree (v. 1.4.2) and annotated in Adobe Photoshop CS6 (Rambaut, 2009).

## Results

### Fungal alpha diversity differs between sample types

Alpha diversity was significantly different between bulk sample types (K–W test; *p* < 0.01, Figure S1, Table S5) for all diversity metrics. Post-hoc Dunn tests identified that the alpha diversity for bulk root and rhizome tissues was consistently lower than the alpha diversity of the sediment (*p* < 0.05, Table S6). This is consistent with previous sequence-based studies of seagrass associated fungi (Hurtado-McCormick et al., 2019). The alpha diversity of the *Z. marina* mycobiome was found to be much lower than that previously seen for the bacterial communities associated with *Z. marina* (Ettinger et al., 2017).

Alpha diversity was not significantly different between root epiphytes and endophytes for any metrics (K-W test, *p* > 0.05, Figure S2). However, alpha diversity was significantly different between leaf epiphytes and endophytes when using the Shannon diversity index which incorporates evenness (K-W test, *p* < 0.05), but was not significantly different when looking at the number of observed ASVs (*p* > 0.05).

### Fungal community structure differs between and within plant parts

Fungal community structure differed significantly between different bulk sample types (PERMANOVA, *p* < 0.001, Table S7, Figure S3). Subsequent pairwise PERMANOVA test results found that community structure was significantly different for all sample type comparisons (*p* < 0.05, Table S8) except sediment vs. roots and leaves vs. roots (*p* > 0.05).

We note, however, that within group variance, also known as dispersion, also differed significantly between bulk sample types (betadisper, *p* < 0.01, Table S9), with rhizomes having more dispersion than sediment or roots (Tukey HSD, *p* < 0.01, Table S10). PERMANOVA results can be confounded by dispersion differences when not using a balanced design with equal sample numbers for categories being compared. In such instances, a significant PERMANOVA result is unable to distinguish between differences in mean dispersions and differences in mean centroids. Thus, these results must be interpreted with circumspection.

Fungal community structure also differed significantly between leaf epiphyte and endophyte communities (PERMANOVA, *p* < 0.001, Table S11, Figure 1) and between leaf length segments (*p* < 0.05). Pair-wise PERMANOVA test results found that the first five inches of leaves had significantly different community structure from the last fifteen inches (*p* < 0.01, Table S12). No significant dispersion differences were detected between the community structure of epiphytes and endophytes (betadisper, *p* > 0.05, Table S13) and the differences in mean dispersions between leaf length segments was barely non-significant (*p* = 0.052). The first five inches of leaves had dispersions that differed significantly from the last ten inches (Tukey HSD, *p* < 0.05, Table S14).

**Figure 1:**
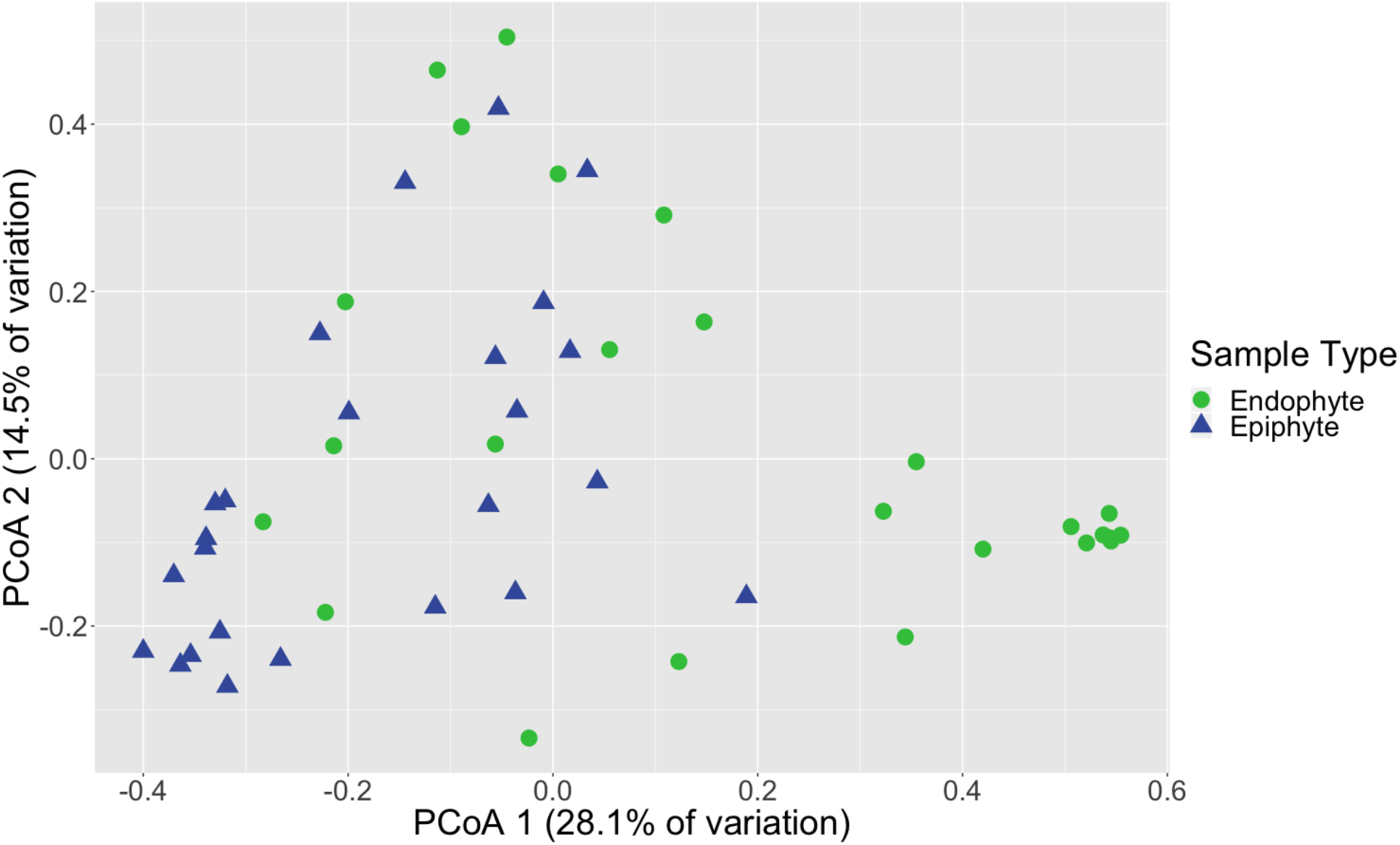
Leaf fungal community structure varies between epiphytes and endophytes. Principal Coordinates Analysis (PCoA) visualization of Bray-Curtis dissimilarities of fungal communities associated with leaves. Points in the ordination are colored and represented by shapes based on epiphyte (blue triangles; n = 25) or endophyte (green circles; n = 25) status. The dataset was first subset to a depth of 5,000 ITS2 amplicon sequences per sample and then Bray-Curtis dissimilarities were calculated using the ordinate function in phyloseq (McMurdie and Holmes, 2013).

### Taxonomic composition of mycobiome

The composition of the mycobiome of bulk samples types was generally comprised of members of the taxonomic orders *Pleosporales, Helotiales, Saccharomycetales, Coniochaetales, Glomerellales, Agaricales, Cystobasidiales* and *Malasseziales* (Table S15, Figure 2). However, we note that the mycobiome of seagrass tissues was dominated by ASVs that were unable to be classified to a specific fungal phylum using current databases.

**Figure 2:**
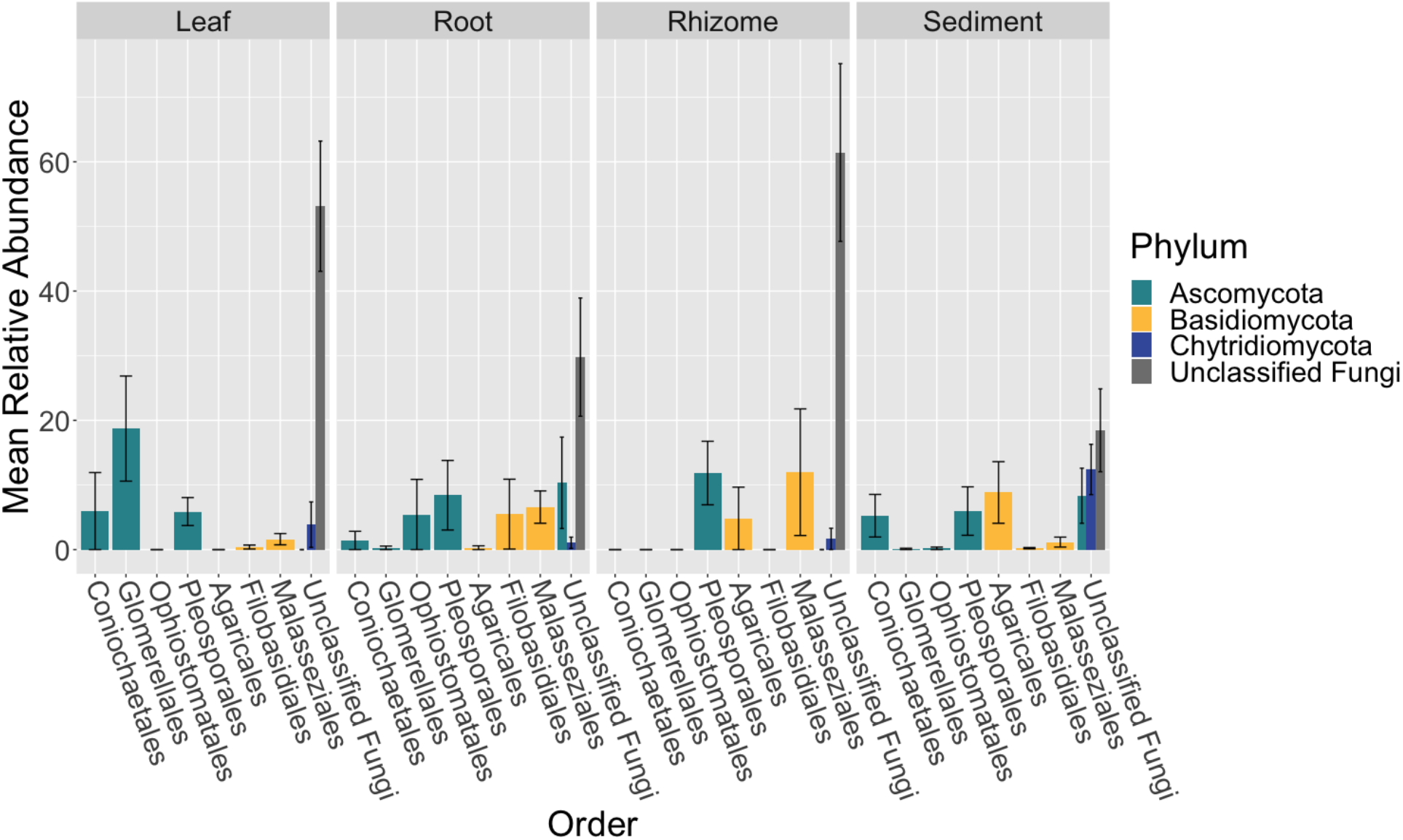
Fungal community composition differs between tissue types. The mean relative abundance of taxonomic orders with a variance of greater than one percent are shown across bulk samples types, leaf (n = 13), root (n = 14), rhizome (n = 7) and sediment (n = 15), with the standard error of the mean represented by error bars and bars colored by taxonomic phylum. The dataset was first subset to a depth of 10,000 sequences per sample, collapsed at the order level using the tax_glom function in phyloseq (McMurdie and Holmes, 2013) and then converted into relative abundance values. Taxonomy was inferred for ITS2 amplicon sequence variants using the RDP Naive Bayesian Classifier algorithm with a modified UNITE (v. 8.0) database (UNITE Community, 2019; Wang et al., 2007).

The order *Agaricales* had a mean relative abundance that was significantly different between sample types (K-W test, *p* < 0.05, Table S16), while the orders *Malasseziales* and *Glomerellales* had mean relative abundances that were only marginally nonsignificant (p = 0.06). *Agaricales* was enriched in the sediment relative to root and leaf tissues (Dunn, *p* < 0.01, Table S17). Whereas *Malasseziales* had a higher mean relative abundance on the roots relative to the leaves and sediment (*p* < 0.01) and *Glomerellales* had an increased abundance on the leaves relative to all other sample types (p < 0.05).

### Mycobiome variation in seagrass leaves

There were no taxonomic orders with a mean relative abundance that was significantly different between leaf segments (K-W test, *p* > 0.05, Table S18, Table S19, Figure 3). However given the relatively high abundance of ASVs that were unable to be assigned confidently to a fungal order, we investigated whether the mean relative abundance of specific ASVs varied across leaf segments (Table S20, Figure 4). We found that the ASV, SV12, had a significantly higher abundance in leaf segments further from the sediment (K-W test, *p* < 0.001, Table S21). This was further supported by post-hoc Dunn test results (*p* < 0.01, Table S22). Based on megablast searches at NCBI, SV12 most closely matches existing ITS2 data from fungal sequences identified as coming from Aphelidomycota. In addition, we were interested in if there were ASVs that had mean relative abundances that varied with epiphyte and endophyte status. We found that the SV8 complex was enriched in epiphyte washes relative to endophyte samples (K-W test, *p* < 0.01, Table S23, Figure S4).

**Figure 3:**
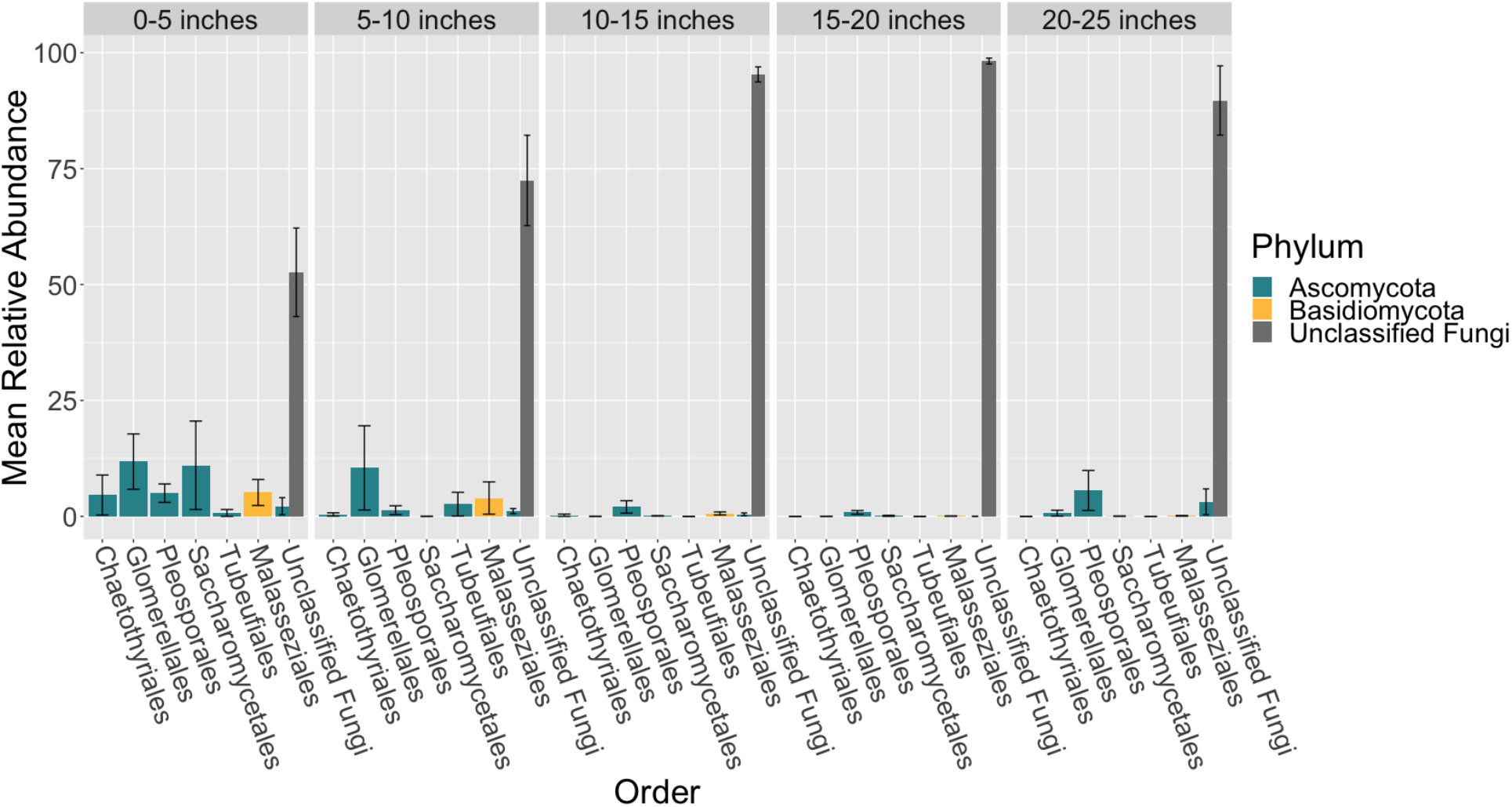
Mean relative abundance of unclassifiable fungi increases across leaf length. The mean relative abundance of taxonomic orders with a variance of greater than 0.1 percent are shown across leaf segments, 0-5 inches (n = 10), 5-10 inches (n = 10), 10-15 inches (n = 10), 15-20 inches (n = 10) and 20-25 inches (n = 10), with the standard error of the mean represented by error bars and bars colored by taxonomic phylum. The dataset was first subset to a depth of 5,000 sequences per sample, collapsed at the order level using the tax_glom function in phyloseq (McMurdie and Holmes, 2013) and then converted into relative abundance values. Taxonomy was inferred for ITS2 amplicon sequence variants using the RDP Naive Bayesian Classifier algorithm with a modified UNITE (v. 8.0) database (UNITE Community, 2019; Wang et al., 2007).

**Figure 4:**
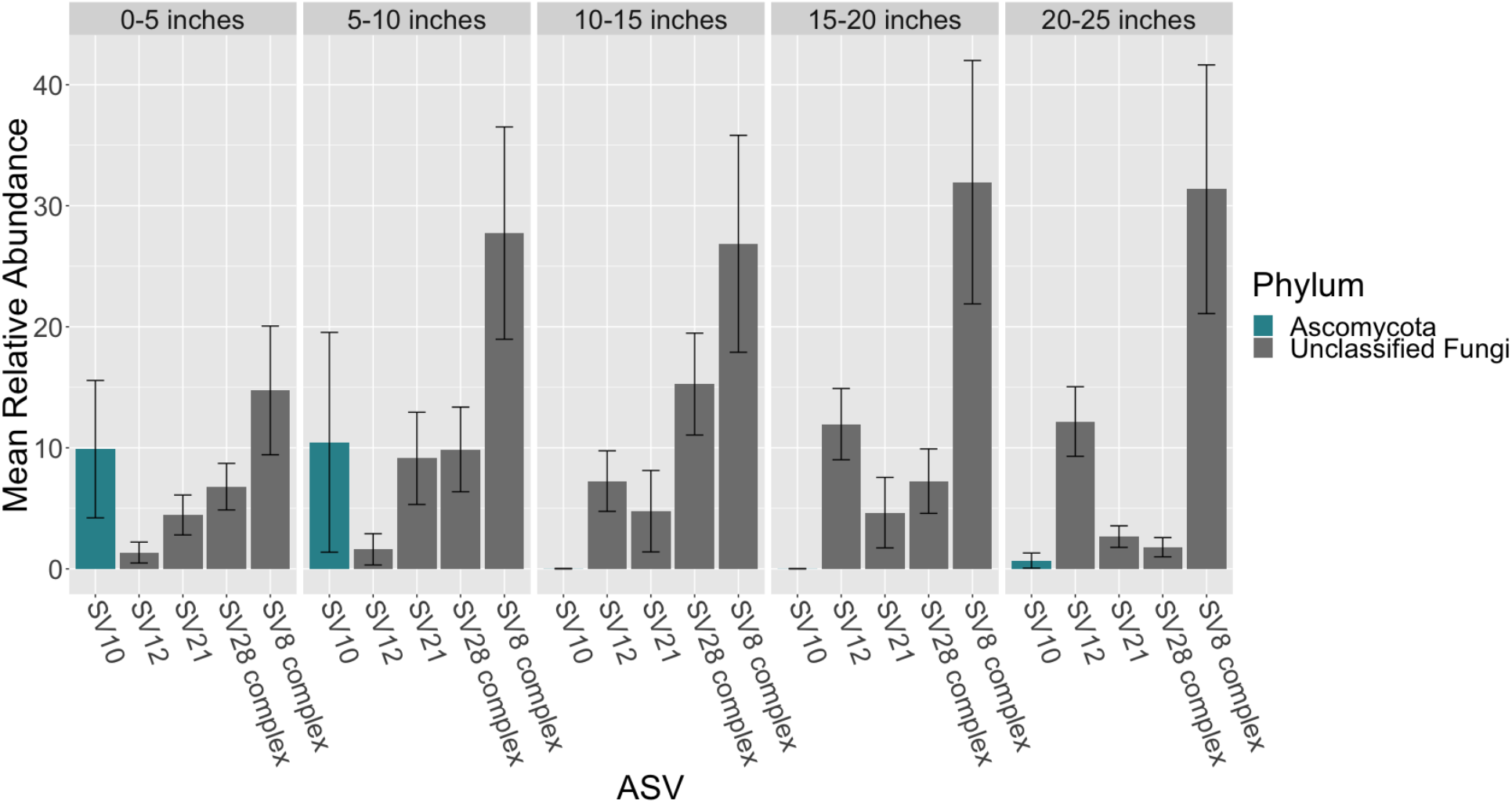
Mean relative abundance of amplicon sequence variants (ASVs) across leaf length. The mean relative abundance of ASVs with a mean of greater than two percent are shown across leaf segments, 0-5 inches (n = 10), 5-10 inches (n = 10), 10-15 inches (n = 10), 15-20 inches (n = 10) and 20-25 inches (n = 10), with the standard error of the mean represented by error bars and bars colored by taxonomic phylum. The dataset was first subset to a depth of 5,000 sequences per sample and then converted into relative abundance values. ASVs were grouped into complexes if ASVs shared greater than 99% sequence identity. Taxonomy was inferred for ITS2 ASVs using the RDP Naive Bayesian Classifier algorithm with a modified UNITE (v. 8.0) database (UNITE Community, 2019; Wang et al., 2007).

### Phylogenetic placement and identification of the SV8 complex

Given the high relative abundance of the SV8 complex in the leaves (Figure 4) and its variable status between epiphyte and endophyte samples, we sought to obtain the sequence of the 28S rRNA gene region for the organism matching this ASV. We designed a primer specific to the ITS2 ASV for the complex using the sequence of SV8 and used a universal fungal 28S rRNA gene primer to obtain a linked ITS2-28S rRNA gene sequence associated with this complex. The resulting ITS2 portion of the resulting PCR products align well with all ASVs from the SV8 complex. Therefore, we believe the 28S rRNA gene sequences obtained accurately represent the flanking region to the taxonomic group represented by the SV8 complex. We then used these 28S rRNA gene sequences to place the organism represented by this ASV complex into a phylogenetic context and assign it a taxonomic classification. This analysis revealed that the SV8 complex (and thus also all ASVs within this complex) groups within the order Lobulomycetales in the phylum Chytridiomycota (Figure 5). Specifically, the SV8 complex is nested within the recently defined Novel Clade SW-I (Van den Wyngaert et al., 2018) and is sister to several culture independent sequences obtained from the marine ecosystem (Hassett et al., 2017).

**Figure 5:**
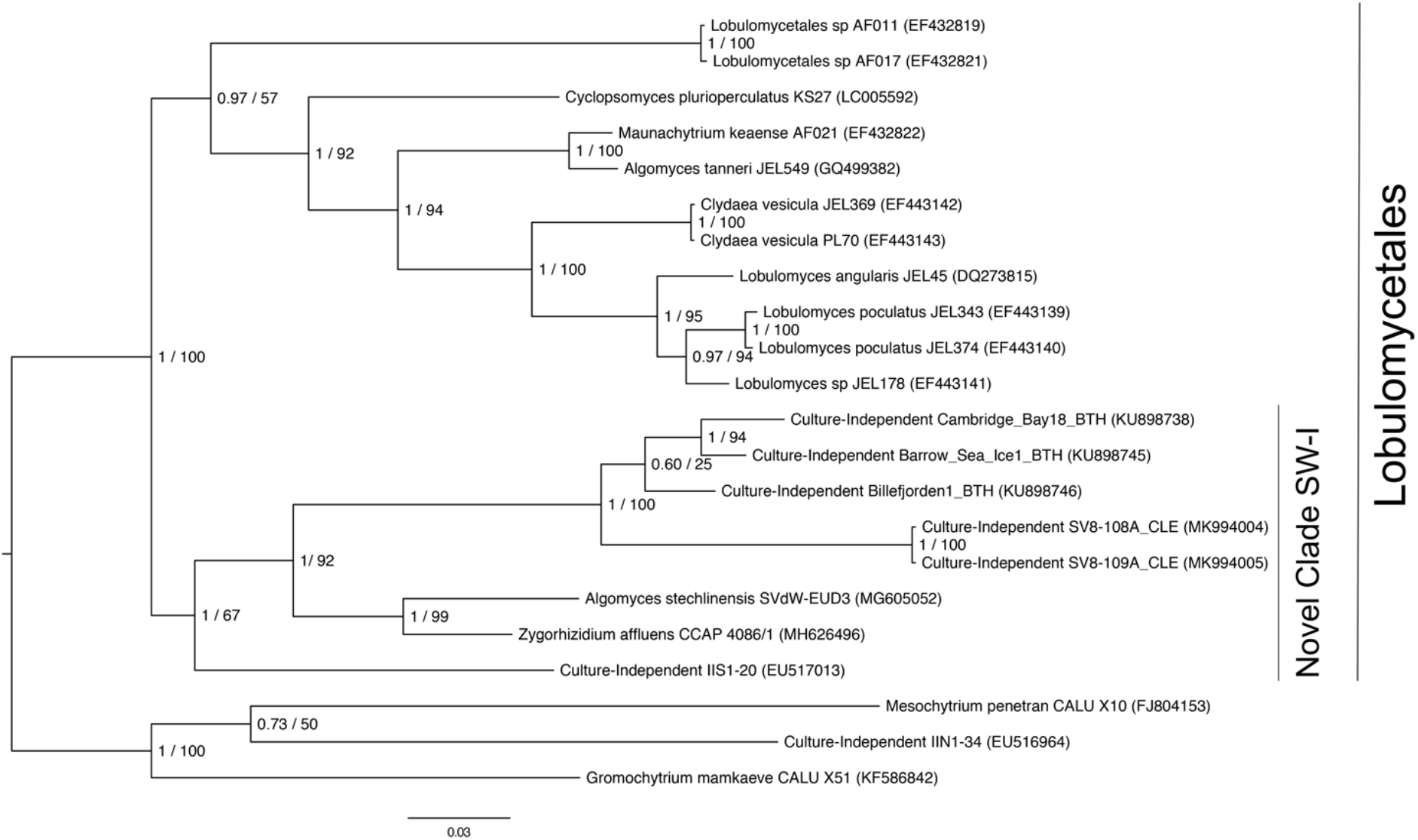
Phylogenetic placement of the SV8 complex in Chytridiomycota. A molecular phylogeny of 28S rRNA genes was constructed using Bayesian inference. Sequence alignments were generated using MAFFT (v. 7.402) on the CIPRES Science Gateway web server, trimmed using trimAl (v.1.2) and a phylogenetic tree was inferred on the trimmed alignment with a GTR + G + I model using MrBayes (v. 3.2.2) (Huelsenbeck and Ronquist, 2001; Katoh, 2002; Miller et al., 2010). Displayed at each node in the tree are the Bayesian posterior probabilities (first number) estimated with MrBayes (v. 3.2.2) and the maximum likelihood bootstrap values (second number) calculated with RAxML (v. 8) (Stamatakis, 2014). The SV8 complex represents a series of four abundant ASVs (SV8, SV11, SV16, SV56) that are greater than 99% identical. The 28S rRNA gene sequences for the SV8 complex are named SV8-108A_CLE and SV8-109A_CLE in this phylogeny and were amplified from two leaf epiphyte samples (sample IDs: 108A and 109A). The GenBank accession numbers of the sequences used to build this phylogeny can be found in Table S4.

## Discussion

Here we offer an in depth survey of the fungi associated with *Z. marina*. We observed that the mycobiome of *Z. marina* roots and rhizomes had lower alpha diversity than that of rhizosphere sediment and that generally the *Z. marina* mycobiome had relatively low species diversity. Low levels of fungal colonization as well as decreased alpha diversity compared to nearby sediment have previously been made in culture and clone based studies of seagrasses (Devarajan and Suryanarayanan, 2002; Ling et al., 2015; Sakayaroj et al., 2010; Shoemaker and Wyllie-Echeverria, 2013; Van den Wyngaert et al., 2018; Venkatachalam et al., 2015). Possible reasons for low abundance and/or infrequent colonization of fungi include indirect factors such as the high salinity and low oxygen levels that are reflective of the marine ecosystem (Nielsen et al., 1999) and direct factors such as seagrass-derived phenolic compounds which are known to act as antimicrobials (Zapata and McMillan, 1979). Additionally, seasonal differences in fungal colonization of seagrasses have been observed, adding an additional layer of complexity and thus making fungal diversity and abundance comparisons difficult (Mata and Cebrián, 2013).

Previous sequence-based studies of seagrass associated fungi, observed high abundances of sequences classified as Pleosporales (Hurtado-McCormick et al., 2019) or Eurotiales (Wainwright et al., 2018), where as here the dominant community members appear to be from understudied lineages like the Chytridiomycota. Additionally, Hurtado-McCormick et al. found evidence of Glomeromycota (arbuscular mycorrhizal fungi) in their dataset, whereas we did not. Possible reasons for these different observations include that these studies sampled different seagrass species than we did, had different sampling schema and used different sequencing methods and primer sets. The use of different sequencing primers specifically has been found to have drastic effects on the results and conclusions of mycobiome studies (Frau et al., 2019).

We did see evidence of known DSE associating with *Z. marina*, including members of the *Pleosporales* and *Glomerellales*, both of which include groups that are known to associate with terrestrial plants (Arnold et al., 2007). We found that the *Glomerellales*, particularly a *Colleotrichum* species (SV10), was abundant in the lower 10 inches of leaf tissue and enriched on the leaves relative to other sample types. DSE, particularly members of the Pleosporales, have previously been observed associating with seagrasses (Borovec and Vohník, 2018; Gnavi et al., 2014; Panno et al., 2013; Torta et al., 2015; Vohník et al., 2015, 2016, 2017, 2019). In the mediteranean seagrass, *Posidonea*, it has been observed that the abundance of the dominant root associated fungi, a Pleosporales species (*Posidoniomyces atricolor* gen. et sp. nov.), is associated with changes in root hair morphology and can form ecto-mycorrhizal-like structures suggesting a close seagrass-fungi symbiosis (Borovec and Vohník, 2018; Vohník et al., 2016, 2017, 2019).

Although many of the ASVs observed here associated with *Z. marina* are still unidentified, we were able to phylogenetically place the SV8 complex in Novel Clade SW-I in the Lobulomycetales. Known members of Novel Clade SW-I are parasitic chytrids of freshwater algae and diatoms (Frenken et al., 2017; Rad-Menéndez et al., 2018; Van den Wyngaert et al., 2018). It is possible that the SV8 complex’s high relative abundance both on and in *Z. marina* leaves may indicate that it is directly infecting host plant tissues and suggests that seagrasses may provide a novel niche for marine chytrids. However, we note that there is another possible interpretation, which is that the SV8 complex is infecting algae or diatoms that are tightly associated with the leaves of *Z. marina* which are themselves a part of the larger *Z. marina* microbiome. This latter interpretation seems to be supported by the observation that the SV8 complex has a higher mean relative abundance in epiphyte washes relative to endophyte washes. However, the closest cultivated relative of the SV8 complex forms sporangia outside of host cells (Van den Wyngaert et al., 2018) and thus, we might expect to find more zoospores in the epiphyte washes. Thus the higher relative abundance of the SV8 complex in the epiphyte washes is consistent with both explanations.

Our observations here are not the first evidence of chytrid associations with seagrasses. A rhizomycelial chytrid was previously observed as the most abundant fungus associating with *Thalassia testudinum* leaves (Newell and Fell, 1980) and was hypothesized to be a possible symbiont, or weak parasite, of the leaves. In a later study, the authors looked for this same chytrid association in *Zostera* with little success (Newell, 1981). However the authors only looked at one location at one time and we note that the SV8 complex was not prevalent during our first two sampling time points. The high abundance of the SV8 complex during our third sampling time point may be because we observed it during a bloom event. Members of the Lobulomycetales have previously been seen to bloom in late summer, which is when we sampled (Frenken et al., 2017).

Another ASV of interest, which was initially unable to be classified into a fungal phylum, is SV12, which most closely matches existing ITS2 data from fungal sequences in the Aphelidomycota. Known Aphelidomycota are intracellular parasites of green algae (Tedersoo et al., 2018). The increased relative abundance of SV12 in the endophyte samples relative to the epiphyte washes is consistent with the possibility that the SV12 containing organisms are directly infecting *Z. marina* leaf tissue. However, as with the SV8 complex, it is also possible that SV12 containing organisms could be associating with an algae that is itself tightly associated with *Z. marina* leaves.

The prevalence of unclassifiable ASVs, many of which have closest matches in the Chytridiomycota and Aphelidomycota, with *Z. marina* makes sense given that these lineages have been previously observed to be the dominant fungal lineages in the marine and aquatic ecosystems (Comeau et al., 2016; Grossart et al., 2019; Picard, 2017; Richards et al., 2012, 2015; Rojas-Jimenez et al., 2019) and have life histories that include associations with green algae (Letcher et al., 2013; Picard et al., 2013; Tedersoo et al., 2018). Thus, it is likely that seagrasses, as marine plants, may be providing these fungal lineages with a new ecological niche.

## Conclusions

We observed that *Z. marina* tissues harbor distinct fungal communities and present analyses and speculation regarding the identity and possible functional roles of these species. For example, we identified that the SV8 complex, which represents the most prevalent sequences associated with *Z. marina* leaf tissue, is nested within Novel Clade SW-I in the order Lobulomycetales and hypothesize that this chytrid may be directly infecting *Z. marina* leaf tissues. However, despite their abundance and possible importance, as evidenced by the number of novel fungal sequences observed here, we still no relatively little about the diversity and functional roles of marine fungi in the seagrass ecosystem. We propose that seagrass beds may be hotspots of marine fungal diversity, specifically the diversity of understudied lineages that are distantly related from well studied fungi (e.g. the Dikarya). Additionally, seagrasses may be an interesting model with which to look at terrestrial to marine transitions in the context of host-microbe interactions since their evolutionary history (coevolution from a terrestrial ancestor) mirrors the evolutionary history of many lineages of marine fungi, which are thought to have ancestors that secondarily returned to the marine environment multiple times (Jones et al., 2015; Schoch et al., 2009; Spatafora et al., 1998; Suetrong et al., 2009). This work helps lay a foundation for future seagrass-fungal studies and highlights a need for future studies focusing on marine fungi and the potential functional importance of these understudied communities to the larger seagrass ecosystem.

## Supporting information

Supplemental RMarkdown file

## Author Contributions

Cassandra L. Ettinger conceived and designed the experiments, performed sampling, analyzed the data, prepared figures and/or tables, wrote and reviewed drafts of the paper. Jonathan A. Eisen advised on data analysis, edited and reviewed drafts of the paper.

## DNA Deposition

This ITS amplicon sequencing project has been deposited at GenBank under accession no. <u>PRJNA515720</u> and consensus sequences for the SV8 complex were deposited under accession no. <u>MK994004, MK994005</u>.

## Acknowledgements

We would like to thank Katherine Dynarski (ORCID: <u>0000-0001-5101-9666</u>) and Sonia Ghose (ORCID: <u>0000-0001-5667-6876</u>) for help with sample collection. We would like to thank John J. Stachowicz for use of his scientific sampling permit, California Department of Fish and Wildlife Scientific Collecting Permit # SC 4874. We also are grateful to Marina LaForgia (ORCID: 0000-0003-4377-0841) for her helpful comments on the R code used.

