## Supplemental Figures & Table Legends for "Characterization of the mycobiome of the seagrass, *Zostera marina*, reveals putative associations with marine chytrids"

**Supplemental Table Legends:**

**Table S1: Number of samples extracted for bulk sample type comparisons**
The number of bulk sample types (leaf, root, rhizome, sediment) for which DNA was extracted listed by location (Westside Point, Gaffney Point) and timepoint (T1, T2, T3). Control sediment was collected from outside the seagrass bed.

**Table S2: Number of samples extracted for intra-plant and epiphyte / endophyte analysis**
The number of samples (leaf, root, rhizome, sediment) for which DNA was extracted for intra-plant (across leaf length and sediment depth) analysis. The leaf and root samples described here were also used to generate epiphyte and endophyte data.

**Table S3: Number of control samples extracted during this experiment**
The number of positive (Zymo Mock Community) and negative (rinse water, ethanol, epiphyte buffer and kit controls) for which DNA was extracted during this experiment.

**Table S4: Sequences used in SV8 complex molecular phylogeny**
Information about the 28S rRNA gene sequences used to generate Figure 5, including the clade, species, strain ID, GenBank Accession number and study of origin for each sequence used.

**Table S5: Kruskal–Wallis tests on alpha diversity metrics**
We used Kruskal–Wallis tests to assess whether alpha diversity was significantly different between categories. We used two different measurements of alpha diversity, observed number of ASVs and the Shannon Diversity index. Categories examined included bulk sample type (leaf, root, rhizome, sediment), root epiphyte / endophyte status and leaf epiphyte / endophyte status.

**Table S6: Post-hoc Dunn tests assessing alpha diversity between bulk sample types**
Alpha diversity was found to be significantly different across bulk sample types (Table S5). We then used a post-hoc Dunn test to examine which sample type comparisons were stochastically dominant on two measurements of alpha diversity, observed number of ASVs and the Shannon Diversity index.

**Table S7: Permanova results of beta diversity of bulk sample types**
PERMANOVA tests were performed to find significant differences in fungal beta diversity, calculated as the Bray Curtis dissimilarity metric, between different categorical variables including sample type (leaf, root, rhizome, sediment), timepoint (T1, T2, T3), replicate core number (1-5), sample location in 96-well sequencing plates sent to Zymo Research, Inc, randomized DNA extraction group (A-P), and DNA extraction kit lot number.

**Table S8: Pairwise PERMANOVA results of beta diversity of bulk sample types**

Here we compared fungal community structure, calculated as Bray Curtis, between pairwise sample types to assess between which sample types communities differed significantly.

**Table S9: Betadisper results of beta diversity of bulk sample types**

Betadisper was used to look for significant differences in the dispersions of different categorical variables when investigating fungal beta diversity, calculated via the Bray Curtis dissimilarity metric. Categorical variables tested included sample type (leaf, root, rhizome, sediment), timepoint (T1, T2, T3), replicate core number (1-5), sample location in 96-well sequencing plates sent to Zymo Research, Inc, randomized DNA extraction group (A-P), and DNA extraction kit lot number.

**Table S10: Tukey HSD Post-hoc tests of beta diversity dispersions**

Tukey HSD post-hoc tests were used to further investigate pairwise combinations for categories with dispersions that differed significantly ($p < 0.05$) as indicated by the Betadisper results (Table S9). These categories included sample type (leaf, root, rhizome, sediment) and randomized DNA extraction group (A-P).

**Table S11: Permanova results of beta diversity of leaf epiphytes and endophytes**

PERMANOVA tests were performed to find significant differences in fungal beta diversity, calculated as the Bray Curtis dissimilarity metric, between different categorical variables including epiphyte / endophyte status, leaf length segment (0-5 inches, 5-10 inches, 10-15 inches, 15-20 inches, 20-25 inches), replicate core number (1-5), sample location in 96-well sequencing plates sent to Zymo Research, Inc, randomized DNA extraction group (A-P), and DNA extraction kit lot number.

**Table S12: Pairwise PERMANOVA results of beta diversity of leaf length**

Here we compared fungal community structure, calculated as Bray Curtis, between pairwise leaf length segments to assess between which segments had community structures that differed significantly.

**Table S13: Betadisper results of beta diversity of leaf epiphytes and endophytes**

Betadisper was used to look for significant differences in the dispersions of different categorical variables when investigating fungal beta diversity, calculated via the Bray Curtis dissimilarity metric. Categorical variables tested included epiphyte / endophyte status, leaf length segment (0-5 inches, 5-10 inches, 10-15 inches, 15-20 inches, 20-25 inches), replicate core number (1-5), sample location in 96-well sequencing plates sent to Zymo Research, Inc, randomized DNA extraction group (A-P), and DNA extraction kit lot number.

**Table S14: Tukey HSD Post-hoc tests of beta diversity dispersions**

Tukey HSD post-hoc tests were used to further investigate pairwise combinations of leaf length segments since segments were found to have dispersions that differed significantly ($p < 0.05$) as indicated by the Betadisper results (Table S13).

**Table S15: Mean, standard deviation and standard error of the relative abundances of taxonomic orders across bulk sample types**

Only orders with a mean relative abundance with a variance of greater than or equal to one percent are included here.

**Table S16: Kruskal-Wallis tests of mean relative abundance of taxonomic orders across bulk sample types**

We used Kruskal–Wallis tests to assess whether the mean relative abundance of taxonomic orders was significantly different between bulk sample types (leaf, root, rhizome, sediment).

**Table S17: Post-hoc Dunn tests of mean relative abundance of taxonomic orders across bulk sample types**

For taxonomic orders with uncorrected p-values that were significantly different ($p < 0.05$) between bulk sample types (Table S16), we then used post-hoc Dunn test to examine which sample type comparisons were driving these differences.

**Table S18: Mean, standard deviation and standard error of the relative abundances of taxonomic orders across leaf length**

Only orders with a mean relative abundance with a variance of greater than or equal to 0.1 percent are included here.

**Table S19: Kruskal-Wallis tests of mean relative abundance of taxonomic orders across leaf length**

We used Kruskal–Wallis tests to assess whether the mean relative abundance of taxonomic orders was significantly different between leaf length segments (0-5 inches, 5-10 inches, 10-15 inches, 15-20 inches, 20-25 inches).

**Table S20: Mean, standard deviation and standard error of the relative abundances of ASVs across leaf length**

Only ASVs with a mean relative abundance with of greater than or equal to two percent are included here.

**Table S21: Kruskal-Wallis tests of mean relative abundance of ASVs across leaf length**

We used Kruskal–Wallis tests to assess whether the mean relative abundance of ASVs was significantly different between leaf length segments (0-5 inches, 5-10 inches, 10-15 inches, 15-20 inches, 20-25 inches).

**Table S22: Post-hoc Dunn tests of mean relative abundance of ASVs across leaf length**

For ASVs with uncorrected p-values that were significantly different ($p < 0.05$) between bulk sample types (Table S16), we then used post-hoc Dunn test to examine which sample type comparisons were driving these differences.

**Table S23: Kruskal-Wallis tests of mean relative abundance of ASVs between leaf epiphytes and endophytes**

We used Kruskal–Wallis tests to assess whether the mean relative abundance of ASVs was significantly different between leaf epiphyte and endophyte samples.

### Supplemental Figures:

**Figure S1: Alpha diversity does not differ between bulk sample types**

Two alpha diversity metrics, observed number of amplicon sequence variants (ASVs) (left) and Shannon diversity index (right), are depicted as boxplots colored by bulk sample type, leaf (n = 13), root (n = 14), rhizome (n = 7) and sediment (n = 15). The dataset was first subset to a depth of 10,000 sequences per sample and then boxplots were constructed using the plot_richness function in phyloseq.

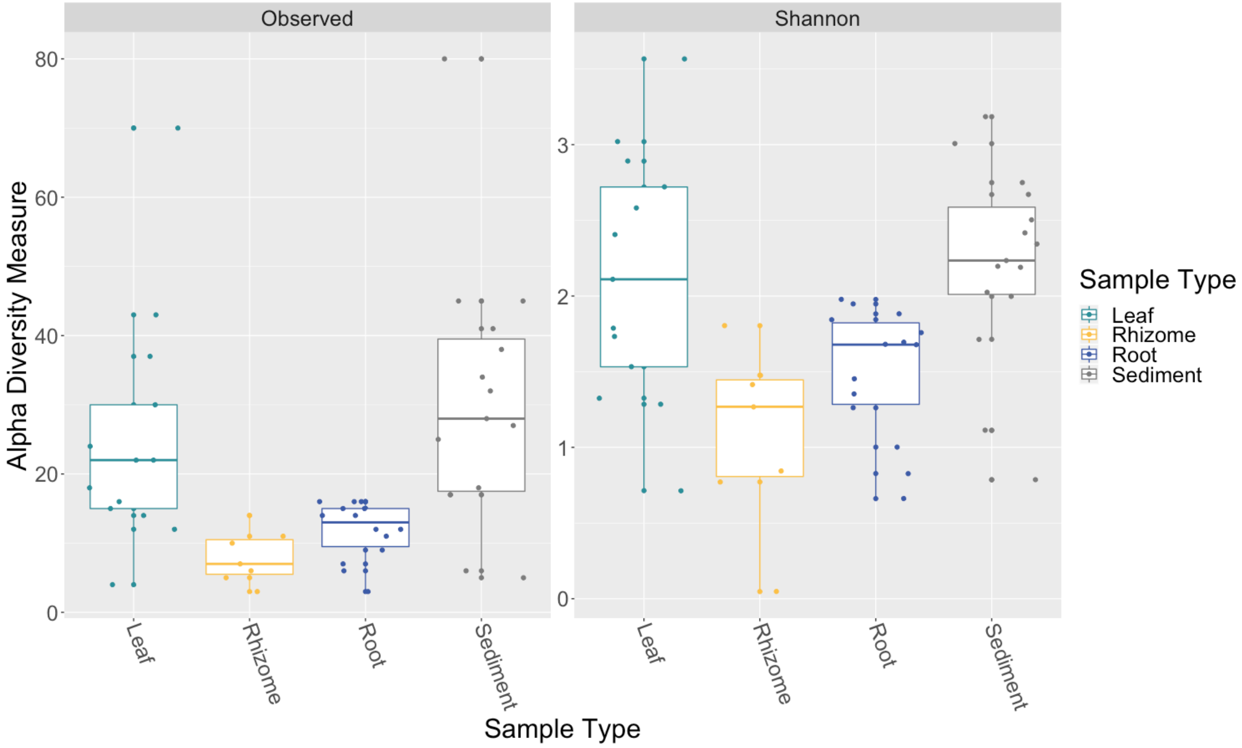

**Figure S2: Alpha diversity differs between leaf epiphyte and endophytes, but not root epiphytes and endophytes**

Two alpha diversity metrics, observed number of amplicon sequence variants (ASVs) (left) and Shannon diversity index (right), are depicted as boxplots split by tissue type, leaf ($n_{total}$ = 50) or root ($n_{total}$ = 35). Samples are further split and colored by epiphyte ($n_{leaf}$ = 25; root, $n_{root}$ = 21) or endophyte ($n_{leaf}$ = 25; $n_{root}$ = 14) status (teal or yellow respectively). The dataset was first subset to a depth of 5,000 sequences per sample and then boxplots were constructed using the plot_richness function in phyloseq.

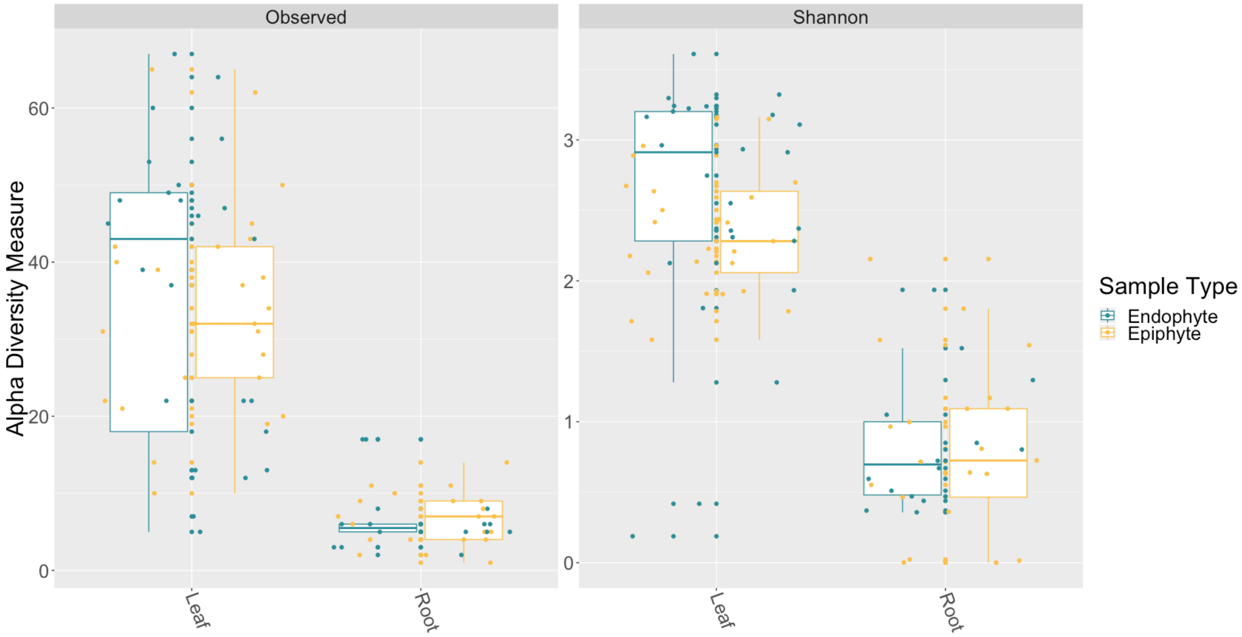

**Figure S3: Fungal community structure varies between bulk sample types**

Principal Coordinates Analysis (PCoA) visualization of Bray-Curtis dissimilarities of fungal communities associated with bulk sample types. Points in the ordination are colored and represented by shapes as follows: leaf (red circle; n = 13), rhizome (green triangle; n = 7), root (blue square; n = 14) and sediment (purple cross; n = 15). The dataset was first subset to a depth of 10,000 ITS2 amplicon sequences per sample and then Bray-Curtis dissimilarities were calculated using the ordinate function in phyloseq.

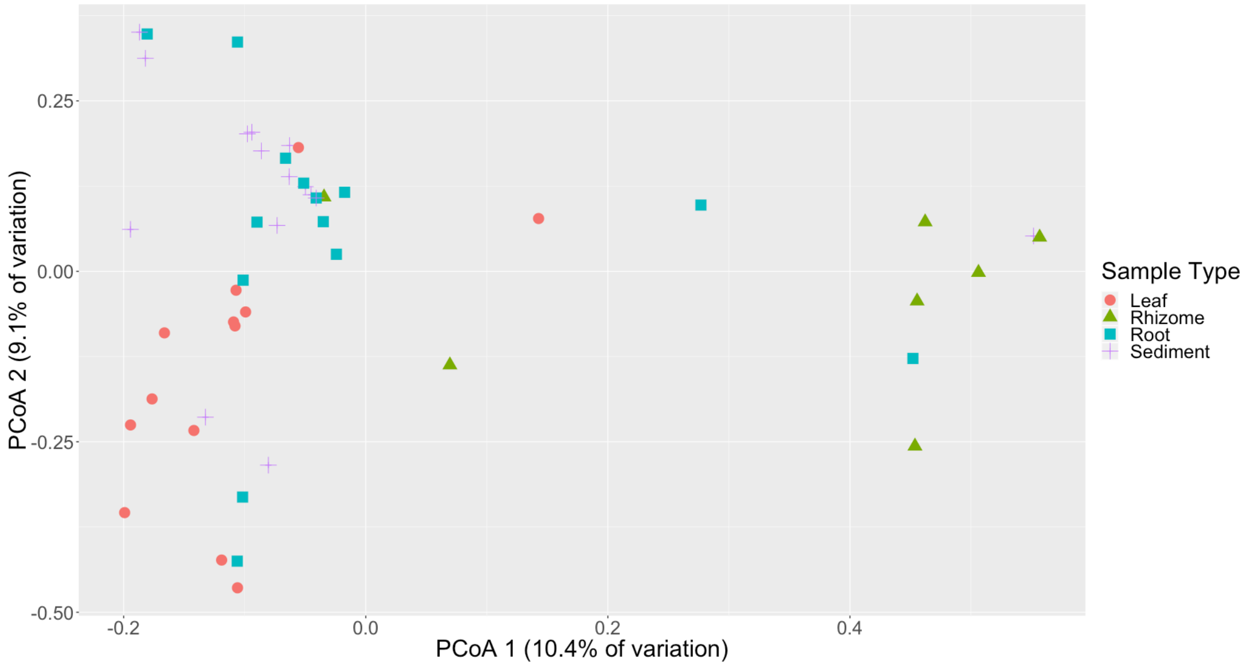

**Figure S4: Differences in mean relative abundance of epiphyte and endophytes ASVs across leaf length**

The mean relative abundance of ASVs with a mean of greater than two percent are shown across leaf segments, 0-5 inches (n = 10), 5-10 inches (n = 10), 10-15 inches (n = 10), 15-20 inches (n = 10) and 20-25 inches (n = 10), with the standard error of the mean represented by error bars and bars colored by taxonomic phylum. Samples are further split in half and the plots are divided up by endophyte (top row) and epiphyte (bottom row) status. The dataset was first subset to a depth of 5,000 sequences per sample and then converted into relative abundance values. ASVs were grouped into complexes if ASVs shared greater than 99% sequence identity. Taxonomy was inferred for ITS2 amplicon sequence variants using the RDP Naive Bayesian Classifier algorithm with a modified UNITE (v. 8.0) database.

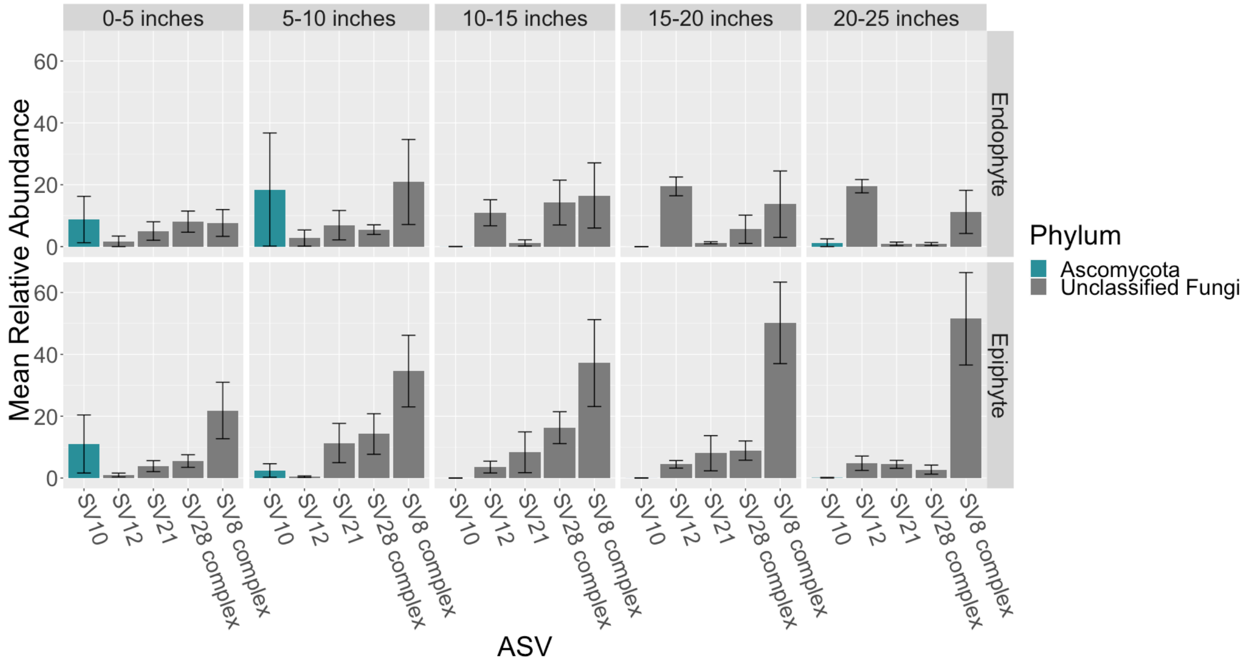
