## Supplemental Tables for "Characterization of the mycobiome of the seagrass, *Zostera marina*, reveals putative associations with marine chytrids"

### ITS2 Amplicon Workflow for Seagrass-associated Fungi Project

*Cassie Ettinger*

#### Project summary

During summer of 2016, I collected cores of seagrass and associated sediment from Bodega Bay. The ITS2 region was amplified and sequenced using DNA from epiphyte washes (root, leaf, rhizome), surface-cleaned tissue samples (root, leaf, rhizome) and sediment.

##### Loading packages and setting up the analysis

First, load in the packages that will be used and note their versions

```
library(ggplot2)
library(qiimer)
library(vegan)
library(ape)
library(Hmisc)
library(phyloseq)
library(RColorBrewer)
library(coin)
library(knitr)
library(rmarkdown)
library(FSA)
library(dplyr)
library(reshape)
library(betapart)
library(dada2)
library(ShortRead)
library(extrafont)
library(tidyverse)
library(magrittr)
library(decontam)

# get versions for the packages, that I am using
# packageVersion('ggplot2') #3.1.0 packageVersion('qiimer')
# packageVersion('vegan') #2.5.5 packageVersion('ape')
# packageVersion('Hmisc') packageVersion('phyloseq') #1.26.0
# packageVersion('RColorBrewer') packageVersion('coin')
# packageVersion('knitr') packageVersion('rmarkdown')
# packageVersion('FSA') #0.8.25 packageVersion('dplyr')
# packageVersion('reshape') packageVersion('betapart')
# packageVersion('dada2') #1.10.0 packageVersion('ShortRead')
# #1.40.0 packageVersion('extrafont')
# packageVersion('tidyverse') packageVersion('magrittr')
# packageVersion('decontam') #1.1.2
```

Going to set the "seed", this ensures any randomization always happens the same way if this analysis needs to be re-run

```
set.seed(5311)
```

##### ITS2 primers used

5.8S_Fun 5' - AACTTTYRRCAAYGGATCWCT - 3'

ITS4_Fun 5' - AGCCTCCGCTTATTGATATGCTTAART - 3'

Primers from:

Taylor, D.L., Walters, W.A., Lennon, N.J., Bochicchio, J., Krohn, A., Caporaso, J.G. and Pennanen, T., 2016. Accurate estimation of fungal diversity and abundance through improved lineage-specific primers optimized for Illumina amplicon sequencing. Applied and environmental microbiology, 82(24), pp.7217-7226.

#### Before using dada2 I removed primers as follows:

```
#Read_ID includes the ID of each read file
for read in $(cat Read_ID.txt);
do cutadapt -g ^AACTTTYRRCAAYGGATCWCT -G ^AGCCTCCGCTTATTGATATGCTTAART
--info-file $read.log.txt --untrimmed-output $read'_R1.untrimmed.fastq'
--untrimmed-paired-output $read'_R2.untrimmed.fastq'
-o $read'_R1.noprimers.fastq' -p $read'_R2.noprimers.fastq'
$read'_R1.fastq' $read'_R2.fastq';
done
```

#### Merging with PEAR

I also merged using PEAR after comparing results to F only reads, R only reads, dada2 merged reads and getting similar results but more reads per sample surviving with PEAR

```
#!/bin/bash -l
#
#SBATCH -n 12 #number cores
#SBATCH -D /share/eisenlab/casett/sgfungi/
#SBATCH -e /share/eisenlab/casett/sgfungi/sgfungi_pear_stderr.txt
#SBATCH -o /share/eisenlab/casett/sgfungi/sgfungi_pear_stderr.txt
#SBATCH -J sgfungi_pear
#SBATCH --mail-type=END # notifications for job done & fail
#SBATCH --mail-user= # send-to address
#SBATCH --mem-per-cpu=4G #memory per node in Gb
#SBATCH -t 96:00:00 #time in hours:min:sec

module load pear
module load perl

#using run_pear.pl from MicrobiomeHelper
#https://github.com/LangilleLab/microbiome_helper
perl /share/eisenlab/casett/sgfungi/scripts/run_pear.pl
-o pear_merged -p 12 /share/eisenlab/casett/sgfungi/*fastq
```

##### Using dada2 to create an amplicon sequence variant (ASV) table

```
# File parsing - For this, we will use only the reads already
# merged with PEAR
path <- "~/Box Sync/Seagrass/SGFungi_RawData/zr1959.rawdata.170922/primers_removed_cutadapt/pear_merged_
filtpath <- file.path(path, "Dada2_Filtered_v1.10.0")
if (!file_test("-d", filtpath)) dir.create(filtpath)
fns <- list.files(path, pattern = "fastq.gz")
```

##### Filter fastqs and store in new directory

```
out <- filterAndTrim(file.path(path, fns), file.path(filtpath,
    fns), maxEE = 2, truncQ = 2, rm.phix = TRUE, compress = TRUE,
    verbose = TRUE, multithread = TRUE)

head(out)
```

##### File parsing of newly filtered files

```
fns = list.files(filtpath)
fns = file.path(filtpath, fns)
filts <- fns[grepl(".fastq.gz$", fns)]
sample.names <- sapply(strsplit(basename(filts), "_"), `[`, 2)
names(filts) <- sample.names
```

##### Get and then inflate errors (to deal with inflated quality score from merging with PEAR)

```
err <- learnErrors(filts, multithread = TRUE, randomize = TRUE,
    nbases = 1e+09)
plotErrors(err, nominalQ = TRUE)

err_3 <- inflateErr(getErrors(err), 3)
plotErrors(err_3, nominalQ = TRUE)
```

##### Sample inference using dada2

```
dds <- vector("list", length(sample.names))
names(dds) <- sample.names
for (sam in sample.names) {
    cat("Processing:", sam, "\n")
    derep <- derepFastq(filts[[sam]])
    dds[[sam]] <- dada(derep, err = err_3, multithread = TRUE)
}

# Construct sequence table
```

```r
seqtab.v5 <- makeSequenceTable(dds)
saveRDS(seqtab.v5, "setab.pear.nomin.err3.dada2.v.1.10.0.rds")
```

##### Remove chimeras

```r
seqtab.nochimera.v5 <- removeBimeraDenovo(seqtab.v5, multithread = TRUE)

saveRDS(seqtab.nochimera.v5, "seqtab.nochimera.pear.nomin.err3.dada.v.1.10.0.rds")
```

##### Fraction of chimeric sequences detected

```r
seqtab.v5 <- readRDS("setab.pear.nomin.err3.dada2.v.1.10.0.rds")
seqtab.nochimera.v5 <- readRDS("seqtab.nochimera.pear.nomin.err3.dada.v.1.10.0.rds")

chimeras = 1 - (sum(seqtab.nochimera.v5)/sum(seqtab.v5))

chimeras
```

```
## [1] 0.003069731
```

##### Inspect sequence length distribution

```r
hist(nchar(getSequences(seqtab.nochimera.v5)))
```

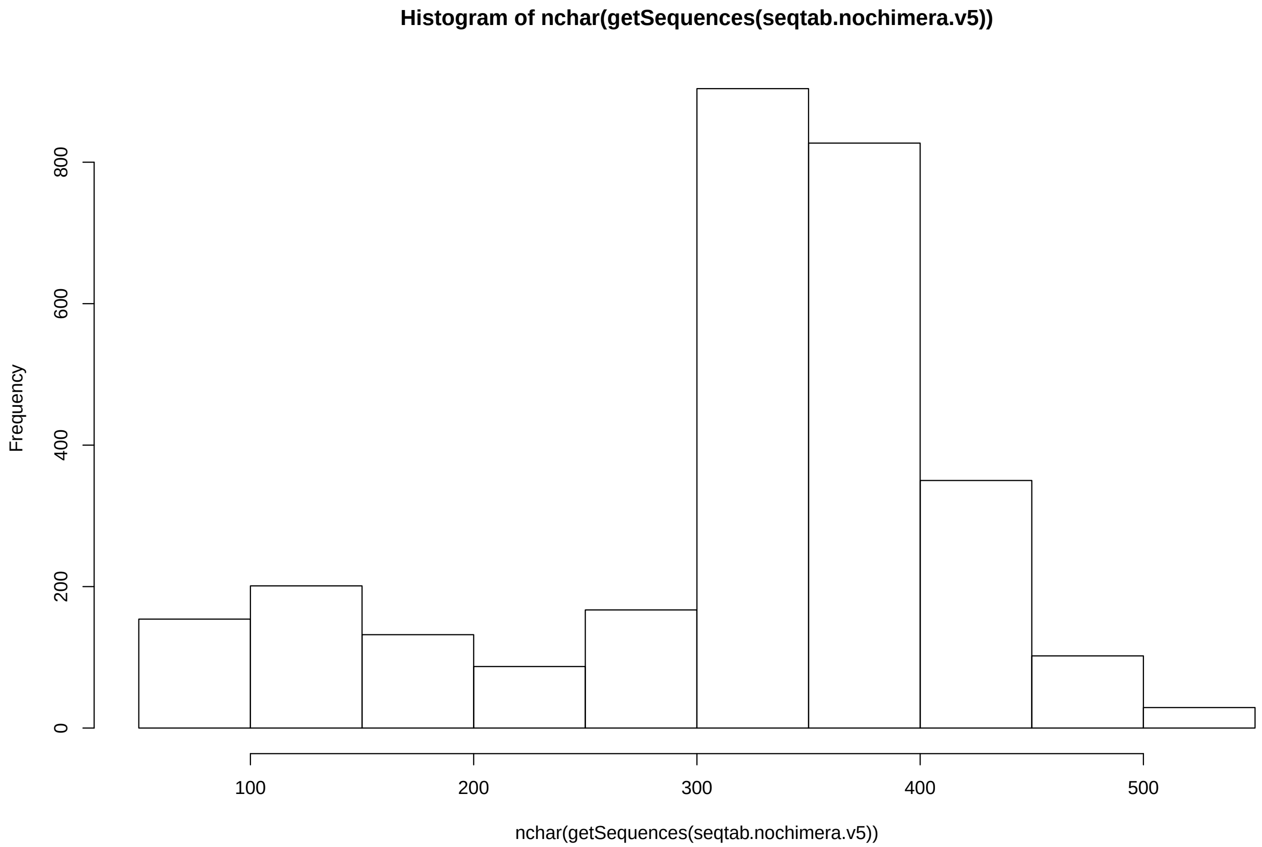


#### Assign Taxonomy using UNITE

```
tax <- assignTaxonomy(seqtab.nochimera.v5, "/Users/Cassie/Box Sync/Databases/unite/sh_general_release_s
    multithread = TRUE, tryRC = TRUE)

saveRDS(tax, "/Users/Cassie/Box Sync/Seagrass/SGFungi_RawData/zr1959.rawdata.170922/primers_removed_cuta
```

#### Get ASVs for ITS-x

```
# Extract sequences from chimera free SV table:
uniquesToFasta(seqtab.nochimera.v5, "unique_ASVs_v5.fasta", ids = paste0("SV",
    seq(length(getSequences(seqtab.nochimera.v5)))))
```

#### Run ITSx on them:

```
./ITSx -i /Users/Cassie/Dropbox/SGFungi/SGFungi R Plots/uniques_ASVs_v5.fasta
-o /Users/Cassie/Dropbox/SGFungi/its_x_results_v5 --date T
--reset T --allow_single_domain --allow_reorder T
--cpu 6 --preserve T --save_regions all
--table T --detailed_results T -t F
```

##### Relabel ASV tables

```
seqtab_final <- seqtab.nochimera.v5
colnames(seqtab_final) <- paste0("SV", 1:ncol(seqtab_final))

tax <- readRDS("/Users/Cassie/Box Sync/Seagrass/SGFungi_RawData/zr1959.rawdata.170922/primers_removed_cu

tax_final <- tax
rownames(tax_final) <- paste0("SV", 1:nrow(tax_final))
```

##### Make phyloseq object

```
mapping <- read.csv("NRS_SG_Fungi_Mapping_File.csv")

otu_table = otu_table(seqtab_final, taxa_are_rows = FALSE)

row.names(mapping) <- paste(row.names(mapping), ".assembled.fastq.gz",
    sep = "")
mapping_file = sample_data(mapping)

taxa_table = tax_table(tax_final)
ps <- phyloseq(otu_table, mapping_file, taxa_table)
```

##### Filter out all sequences that ITS-x did not flag as fungal

```
nf = c("SV7", "SV30", "SV54", "SV72", "SV79", "SV86", "SV94",
    "SV101", "SV103", "SV111", "SV114", "SV118", "SV124", "SV131",
    "SV136", "SV139", "SV146", "SV163", "SV175", "SV205", "SV211",
    "SV223", "SV224", "SV225", "SV254", "SV286", "SV293", "SV312",
    "SV314", "SV315", "SV323", "SV353", "SV362", "SV367", "SV373",
    "SV390", "SV415", "SV448", "SV451", "SV459", "SV464", "SV490",
    "SV491", "SV492", "SV493", "SV498", "SV505", "SV510", "SV512",
    "SV522", "SV550", "SV560", "SV562", "SV575", "SV577", "SV584",
    "SV585", "SV603", "SV613", "SV616", "SV625", "SV649", "SV653",
    "SV660", "SV669", "SV672", "SV680", "SV681", "SV692", "SV710",
    "SV711", "SV712", "SV713", "SV719", "SV721", "SV726", "SV727",
    "SV730", "SV731", "SV740", "SV753", "SV761", "SV762", "SV767",
    "SV776", "SV779", "SV785", "SV797", "SV799", "SV804", "SV814",
    "SV817", "SV823", "SV834", "SV836", "SV843", "SV851", "SV863",
    "SV870", "SV886", "SV887", "SV894", "SV904", "SV921", "SV923",
    "SV941", "SV953", "SV960", "SV961", "SV968", "SV973", "SV977",
    "SV989", "SV992", "SV994", "SV1004", "SV1018", "SV1020",
    "SV1030", "SV1032", "SV1044", "SV1049", "SV1053", "SV1057",
    "SV1067", "SV1078", "SV1087", "SV1095", "SV1100", "SV1106",
    "SV1108", "SV1118", "SV1123", "SV1133", "SV1137", "SV1142",
    "SV1143", "SV1145", "SV1148", "SV1151", "SV1166", "SV1167",
    "SV1183")
nf2 = c("SV1187", "SV1189", "SV1191", "SV1195", "SV1202", "SV1204",
    "SV1212", "SV1227", "SV1229", "SV1230", "SV1249", "SV1250",
    "SV1255", "SV1256", "SV1265", "SV1270", "SV1278", "SV1280",
```

```
    "SV1285", "SV1289", "SV1290", "SV1291", "SV1296", "SV1308",
    "SV1312", "SV1318", "SV1319", "SV1329", "SV1334", "SV1343",
    "SV1346", "SV1355", "SV1358", "SV1361", "SV1364", "SV1367",
    "SV1369", "SV1371", "SV1373", "SV1377", "SV1379", "SV1381",
    "SV1383", "SV1385", "SV1386", "SV1406", "SV1409", "SV1415",
    "SV1419", "SV1427", "SV1428", "SV1431", "SV1432", "SV1437",
    "SV1443", "SV1451", "SV1453", "SV1456", "SV1458", "SV1459",
    "SV1466", "SV1472", "SV1473", "SV1490", "SV1501", "SV1503",
    "SV1504", "SV1509", "SV1512", "SV1526", "SV1528", "SV1530",
    "SV1539", "SV1541", "SV1547", "SV1554", "SV1557", "SV1571",
    "SV1575", "SV1577", "SV1580", "SV1589", "SV1591", "SV1595",
    "SV1596", "SV1597", "SV1599", "SV1605", "SV1608", "SV1615",
    "SV1620", "SV1621", "SV1624", "SV1629", "SV1631", "SV1635",
    "SV1637", "SV1641", "SV1642", "SV1643", "SV1649", "SV1654",
    "SV1655", "SV1656", "SV1658", "SV1659", "SV1669", "SV1676",
    "SV1681", "SV1688", "SV1690", "SV1691", "SV1694", "SV1695",
    "SV1704", "SV1706", "SV1709", "SV1712", "SV1716", "SV1721",
    "SV1723", "SV1724", "SV1725", "SV1726", "SV1729", "SV1731",
    "SV1732", "SV1733", "SV1734", "SV1739", "SV1742", "SV1743",
    "SV1751", "SV1753", "SV1758", "SV1763", "SV1766", "SV1771",
    "SV1773", "SV1774", "SV1781", "SV1784", "SV1787", "SV1791",
    "SV1796", "SV1797", "SV1798", "SV1799", "SV1800", "SV1814",
    "SV1815", "SV1818", "SV1829", "SV1831", "SV1841", "SV1843",
    "SV1847", "SV1849", "SV1852", "SV1861", "SV1862", "SV1865",
    "SV1867", "SV1868", "SV1871", "SV1874", "SV1876", "SV1877",
    "SV1878", "SV1881", "SV1882", "SV1885", "SV1886", "SV1888",
    "SV1895", "SV1897", "SV1899", "SV1902", "SV1903", "SV1904",
    "SV1911", "SV1912", "SV1914", "SV1917", "SV1918", "SV1922",
    "SV1923", "SV1924", "SV1926", "SV1927", "SV1929", "SV1931",
    "SV1932", "SV1934", "SV1944", "SV1946", "SV1950", "SV1953",
    "SV1954", "SV1956", "SV1958", "SV1963", "SV1969", "SV1974",
    "SV1980", "SV1981", "SV1982", "SV1984", "SV1986", "SV1989",
    "SV1990", "SV1992", "SV1997", "SV2000", "SV2002", "SV2003",
    "SV2004", "SV2006", "SV2010", "SV2019", "SV2020", "SV2023",
    "SV2024", "SV2025", "SV2031", "SV2032", "SV2033", "SV2034",
    "SV2035", "SV2036", "SV2037", "SV2040", "SV2042", "SV2044",
    "SV2047", "SV2048", "SV2053", "SV2054", "SV2056", "SV2060",
    "SV2061", "SV2063", "SV2064", "SV2067", "SV2068", "SV2070",
    "SV2071", "SV2072", "SV2074", "SV2075", "SV2077", "SV2079",
    "SV2085", "SV2086", "SV2087", "SV2090", "SV2093", "SV2094",
    "SV2096", "SV2097", "SV2098", "SV2099", "SV2100", "SV2103",
    "SV2107", "SV2108", "SV2111", "SV2114", "SV2117", "SV2118",
    "SV2124", "SV2125", "SV2126", "SV2128", "SV2129", "SV2130",
    "SV2131", "SV2132", "SV2134", "SV2137", "SV2138", "SV2139",
    "SV2140", "SV2141", "SV2142", "SV2143", "SV2144", "SV2149",
    "SV2153", "SV2154", "SV2155", "SV2156", "SV2157", "SV2160",
    "SV2162", "SV2163", "SV2165", "SV2172", "SV2173", "SV2175",
    "SV2176", "SV2177", "SV2179", "SV2180", "SV2182", "SV2183",
    "SV2185", "SV2188", "SV2189", "SV2190", "SV2191", "SV2192",
    "SV2193", "SV2197", "SV2201", "SV2204", "SV2205", "SV2207",
    "SV2208", "SV2210", "SV2211", "SV2212", "SV2214", "SV2215",
    "SV2216", "SV2217", "SV2218", "SV2219", "SV2221", "SV2224",
    "SV2230", "SV2231", "SV2233", "SV2238", "SV2239", "SV2240",
```

```
    "SV2241", "SV2242", "SV2243", "SV2244", "SV2245", "SV2246",
    "SV2247", "SV2248", "SV2251", "SV2254", "SV2258", "SV2261",
    "SV2262", "SV2264", "SV2267", "SV2268", "SV2269", "SV2270",
    "SV2271", "SV2272", "SV2273", "SV2274", "SV2275", "SV2278",
    "SV2280", "SV2281", "SV2283", "SV2284", "SV2285", "SV2286",
    "SV2289", "SV2293", "SV2297", "SV2298", "SV2299", "SV2300",
    "SV2301", "SV2303", "SV2304", "SV2305", "SV2306", "SV2307",
    "SV2308", "SV2309", "SV2310", "SV2311", "SV2312", "SV2314",
    "SV2316", "SV2317", "SV2318", "SV2321", "SV2322", "SV2324",
    "SV2325", "SV2327", "SV2329", "SV2330", "SV2331", "SV2332",
    "SV2333", "SV2334", "SV2335", "SV2336", "SV2337", "SV2338",
    "SV2339")
nf3 = c("SV2340", "SV2341", "SV2342", "SV2343", "SV2344", "SV2345",
    "SV2346", "SV2347", "SV2348", "SV2349", "SV2350", "SV2351",
    "SV2352", "SV2353", "SV2355", "SV2358", "SV2359", "SV2361",
    "SV2364", "SV2365", "SV2369", "SV2373", "SV2376", "SV2378",
    "SV2379", "SV2380", "SV2381", "SV2382", "SV2383", "SV2385",
    "SV2386", "SV2387", "SV2388", "SV2389", "SV2390", "SV2391",
    "SV2392", "SV2393", "SV2394", "SV2395", "SV2396", "SV2397",
    "SV2398", "SV2401", "SV2402", "SV2403", "SV2404", "SV2405",
    "SV2406", "SV2407", "SV2408", "SV2409", "SV2410", "SV2411",
    "SV2412", "SV2413", "SV2414", "SV2416", "SV2420", "SV2421",
    "SV2422", "SV2423", "SV2424", "SV2425", "SV2426", "SV2428",
    "SV2432", "SV2433", "SV2435", "SV2436", "SV2440", "SV2441",
    "SV2443", "SV2444", "SV2445", "SV2446", "SV2447", "SV2448",
    "SV2449", "SV2450", "SV2451", "SV2452", "SV2453", "SV2454",
    "SV2456", "SV2457", "SV2458", "SV2459", "SV2461", "SV2462",
    "SV2463", "SV2464", "SV2465", "SV2466", "SV2467", "SV2468",
    "SV2469", "SV2470", "SV2471", "SV2472", "SV2473", "SV2474",
    "SV2475", "SV2476", "SV2477", "SV2478", "SV2479", "SV2481",
    "SV2482", "SV2483", "SV2484", "SV2485", "SV2486", "SV2487",
    "SV2488", "SV2489", "SV2490", "SV2491", "SV2492", "SV2495",
    "SV2496", "SV2497", "SV2499", "SV2500", "SV2503", "SV2504",
    "SV2509", "SV2513", "SV2514", "SV2516", "SV2517", "SV2518",
    "SV2519", "SV2520", "SV2523", "SV2524", "SV2525", "SV2526",
    "SV2527", "SV2528", "SV2529", "SV2530", "SV2531", "SV2533",
    "SV2534", "SV2535", "SV2536", "SV2537", "SV2538", "SV2539",
    "SV2540", "SV2541", "SV2542", "SV2543", "SV2544", "SV2545",
    "SV2546", "SV2547", "SV2548", "SV2549", "SV2550", "SV2551",
    "SV2552", "SV2553", "SV2554", "SV2555", "SV2556", "SV2557",
    "SV2558", "SV2559", "SV2560", "SV2561", "SV2562", "SV2563",
    "SV2564", "SV2565", "SV2566", "SV2567", "SV2568", "SV2569",
    "SV2570", "SV2571", "SV2572", "SV2573", "SV2574", "SV2575",
    "SV2576", "SV2577", "SV2578", "SV2579", "SV2580", "SV2581",
    "SV2582", "SV2583", "SV2584", "SV2585", "SV2586", "SV2587",
    "SV2588", "SV2589", "SV2590", "SV2591", "SV2592", "SV2593",
    "SV2594", "SV2596", "SV2597", "SV2598", "SV2599", "SV2600",
    "SV2601", "SV2602", "SV2603", "SV2604", "SV2605", "SV2606",
    "SV2607", "SV2608", "SV2609", "SV2610", "SV2611", "SV2612",
    "SV2613", "SV2614", "SV2615", "SV2616", "SV2618", "SV2619",
    "SV2620", "SV2621", "SV2623", "SV2627", "SV2628", "SV2629",
    "SV2630", "SV2631", "SV2632", "SV2634", "SV2635", "SV2636",
    "SV2638", "SV2639", "SV2641", "SV2644", "SV2645", "SV2647",
```

```
    "SV2648", "SV2651", "SV2652", "SV2658", "SV2659", "SV2660",
    "SV2663", "SV2667", "SV2673", "SV2674", "SV2675", "SV2676",
    "SV2677", "SV2679", "SV2680", "SV2681", "SV2682", "SV2683",
    "SV2684", "SV2685", "SV2686", "SV2687", "SV2688", "SV2693",
    "SV2694", "SV2695", "SV2696", "SV2697", "SV2698", "SV2699",
    "SV2700", "SV2701", "SV2702", "SV2703", "SV2704", "SV2705",
    "SV2706", "SV2707", "SV2708", "SV2709", "SV2710", "SV2711",
    "SV2712", "SV2713", "SV2714", "SV2715", "SV2716", "SV2717",
    "SV2718", "SV2719", "SV2720", "SV2721", "SV2722", "SV2723",
    "SV2724", "SV2725", "SV2726", "SV2727", "SV2728", "SV2729",
    "SV2730", "SV2731", "SV2732", "SV2733", "SV2734", "SV2735",
    "SV2736", "SV2737", "SV2738", "SV2739", "SV2740", "SV2742",
    "SV2743", "SV2744", "SV2745", "SV2746", "SV2747", "SV2748",
    "SV2749", "SV2750", "SV2751", "SV2752", "SV2753", "SV2754",
    "SV2755", "SV2756", "SV2757", "SV2758", "SV2759", "SV2760",
    "SV2761", "SV2762", "SV2764", "SV2765", "SV2766", "SV2767",
    "SV2768", "SV2769", "SV2770", "SV2771", "SV2772", "SV2773",
    "SV2774", "SV2775", "SV2776", "SV2777", "SV2778", "SV2779",
    "SV2780", "SV2781", "SV2782", "SV2783", "SV2784", "SV2785",
    "SV2787", "SV2788", "SV2789", "SV2790", "SV2791", "SV2792",
    "SV2793", "SV2794", "SV2795", "SV2796", "SV2797", "SV2798",
    "SV2799", "SV2800", "SV2801", "SV2802", "SV2803", "SV2804",
    "SV2805", "SV2806", "SV2807", "SV2808", "SV2809", "SV2810",
    "SV2811", "SV2812", "SV2813", "SV2814", "SV2815", "SV2816",
    "SV2817", "SV2818")
nf4 = c("SV2819", "SV2820", "SV2821", "SV2822", "SV2823", "SV2824",
    "SV2825", "SV2826", "SV2827", "SV2828", "SV2829", "SV2830",
    "SV2831", "SV2832", "SV2833", "SV2834", "SV2835", "SV2836",
    "SV2837", "SV2838", "SV2839", "SV2840", "SV2841", "SV2842",
    "SV2843", "SV2844", "SV2845", "SV2846", "SV2847", "SV2848",
    "SV2850", "SV2851", "SV2852", "SV2853", "SV2854", "SV2855",
    "SV2856", "SV2857", "SV2858", "SV2859", "SV2860", "SV2861",
    "SV2862", "SV2863", "SV2864", "SV2865", "SV2866", "SV2867",
    "SV2868", "SV2869", "SV2870", "SV2871", "SV2872", "SV2873",
    "SV2874", "SV2875", "SV2876", "SV2877", "SV2878", "SV2879",
    "SV2880", "SV2881", "SV2882", "SV2883", "SV2884", "SV2885",
    "SV2886", "SV2887", "SV2888", "SV2889", "SV2890", "SV2892",
    "SV2893", "SV2894", "SV2895", "SV2896", "SV2897", "SV2898",
    "SV2899", "SV2901", "SV2903", "SV2904", "SV2907", "SV2908",
    "SV2909", "SV2911", "SV2912", "SV2913", "SV2914", "SV2916",
    "SV2917", "SV2918", "SV2919", "SV2920", "SV2921", "SV2922",
    "SV2923", "SV2924", "SV2926", "SV2927", "SV2928", "SV2929",
    "SV2930", "SV2931", "SV2932", "SV2933", "SV2934", "SV2935",
    "SV2936", "SV2937", "SV2938", "SV2939", "SV2940", "SV2941",
    "SV2942", "SV2943", "SV2944", "SV2945", "SV2946", "SV2947",
    "SV2948", "SV2949", "SV2950", "SV2951", "SV2952", "SV2953")

allTaxa = taxa_names(ps)
fungalTaxa <- allTaxa[!(allTaxa %in% nf)]
ps_onlyfungi_v1 = prune_taxa(fungalTaxa, ps)

allTaxa = taxa_names(ps_onlyfungi_v1)
fungalTaxa <- allTaxa[!(allTaxa %in% nf2)]
```

```r
ps_onlyfungi_v2 = prune_taxa(fungalTaxa, ps_onlyfungi_v1)

allTaxa = taxa_names(ps_onlyfungi_v2)
fungalTaxa <- allTaxa[!(allTaxa %in% nf3)]
ps_onlyfungi_v3 = prune_taxa(fungalTaxa, ps_onlyfungi_v2)

allTaxa = taxa_names(ps_onlyfungi_v3)
fungalTaxa <- allTaxa[!(allTaxa %in% nf4)]
ps_onlyfungi = prune_taxa(fungalTaxa, ps_onlyfungi_v3)

ps
```

```
## phyloseq-class experiment-level object
## otu_table()   OTU Table:         [ 2953 taxa and 235 samples ]
## sample_data() Sample Data:       [ 235 samples by 37 sample variables ]
## tax_table()   Taxonomy Table:    [ 2953 taxa by 7 taxonomic ranks ]
```

```r
ps_onlyfungi
```

```
## phyloseq-class experiment-level object
## otu_table()   OTU Table:         [ 1889 taxa and 235 samples ]
## sample_data() Sample Data:       [ 235 samples by 37 sample variables ]
## tax_table()   Taxonomy Table:    [ 1889 taxa by 7 taxonomic ranks ]
```

##### Identify possible contaminants using decontam

```r
## We have 4 different types of NC h20, etoh, diluted redford
## buffer and kit controls each must be done seperately so
## first we need to subset our data so that we remove the
## controls we AREN'T looking at this is in case of overlap
## between negative controls

ps_pear_h20_err3 <- subset_samples(ps_onlyfungi, SampleType !=
    "Control_Buffer")
ps_pear_h20_err3 <- subset_samples(ps_pear_h20_err3, SampleType !=
    "Control_EtoH")
ps_pear_h20_err3 <- subset_samples(ps_pear_h20_err3, SampleType !=
    "Control_Kit")

ps_pear_buf_err3 <- subset_samples(ps_onlyfungi, SampleType !=
    "Control_H20")
ps_pear_buf_err3 <- subset_samples(ps_pear_buf_err3, SampleType !=
    "Control_EtoH")
ps_pear_buf_err3 <- subset_samples(ps_pear_buf_err3, SampleType !=
    "Control_Kit")

ps_pear_etoh_err3 <- subset_samples(ps_onlyfungi, SampleType !=
    "Control_Buffer")
ps_pear_etoh_err3 <- subset_samples(ps_pear_etoh_err3, SampleType !=
    "Control_Kit")
ps_pear_etoh_err3 <- subset_samples(ps_pear_etoh_err3, SampleType !=
    "Control_H20")
```

```
ps_pear_kit_err3 <- subset_samples(ps_onlyfungi, SampleType !=
    "Control_Buffer")
ps_pear_kit_err3 <- subset_samples(ps_pear_kit_err3, SampleType !=
    "Control_EtoH")
ps_pear_kit_err3 <- subset_samples(ps_pear_kit_err3, SampleType !=
    "Control_H20")
```

We do not have input DNA concentration, so we are using decontom's "prevalence" method. In this method, the prevalence (presence/absence across samples) of each sequence feature in true positive samples is compared to the prevalence in negative controls to identify contaminants.

Using a strict threshold (threshold=0.5), which will identify as contaminants all sequences thare are more prevalent in negative controls than in positive samples

```
contamdf.prev.pear.h20.05_err3 <- isContaminant(ps_pear_h20_err3,
    method = "prevalence", neg = "IsControl", threshold = 0.5)
contamdf.prev.pear.etoh.05_err3 <- isContaminant(ps_pear_etoh_err3,
    method = "prevalence", neg = "IsControl", threshold = 0.5)
contamdf.prev.pear.buf.05_err3 <- isContaminant(ps_pear_buf_err3,
    method = "prevalence", neg = "IsControl", threshold = 0.5)
contamdf.prev.pear.kit.05_err3 <- isContaminant(ps_pear_kit_err3,
    method = "prevalence", neg = "IsControl", threshold = 0.5)


table(contamdf.prev.pear.h20.05_err3$contaminant)  #7
table(contamdf.prev.pear.etoh.05_err3$contaminant)  #14
table(contamdf.prev.pear.buf.05_err3$contaminant)  #18
table(contamdf.prev.pear.kit.05_err3$contaminant)  #12

# which ASV is contaminant

which(contamdf.prev.pear.h20.05_err3$contaminant)
# 1 2 14 16 32 95 204
which(contamdf.prev.pear.etoh.05_err3$contaminant)
# 1 3 8 13 26 28 31 49 50 55 95 125 144 426
which(contamdf.prev.pear.buf.05_err3$contaminant)
# 1 2 4 5 13 18 19 22 29 32 59 71 82 86 92 172 361 484
which(contamdf.prev.pear.kit.05_err3$contaminant)
# 1 2 3 8 13 14 32 41 43 56 459 593


contaminants.pear.h20_err3 <- subset(contamdf.prev.pear.h20.05_err3,
    contaminant == "TRUE")
contaminants.pear.etoh_err3 <- subset(contamdf.prev.pear.etoh.05_err3,
    contaminant == "TRUE")
contaminants.pear.buf_err3 <- subset(contamdf.prev.pear.buf.05_err3,
    contaminant == "TRUE")
contaminants.pear.kit_err3 <- subset(contamdf.prev.pear.kit.05_err3,
    contaminant == "TRUE")

# save these files as csv to investigate

write.csv(contaminants.pear.h20_err3, "contaminants.pear.h20_err3_v5_f.csv")
write.csv(contaminants.pear.etoh_err3, "contaminants.pear.etoh_err3_v5_f.csv")
write.csv(contaminants.pear.buf_err3, "contaminants.pear.buf_err3_v5_f.csv")
```

```r
write.csv(contaminants.pear.kit_err3, "contaminants.pear.kit_err3_v5_f.csv")
```

We combined these outputs in Excel (sorry for lack of reproducability!) and removed duplicates

```r
contaminants.pear_err3_v5 <- read.csv("contaminants.pear_err3_v5_combined_noduplicates_f.csv")

contaminants.pear_err3_v5.list <- contaminants.pear_err3_v5$X
length(contaminants.pear_err3_v5.list)  #38 contaminants
```

```
## [1] 38
```

#### We will want to remove these ASVs from further analyses

```r
allTaxa = taxa_names(ps_onlyfungi)
newTaxa <- allTaxa[!(allTaxa %in% contaminants.pear_err3_v5.list)]
ps_onlyfungi_NC = prune_taxa(newTaxa, ps_onlyfungi)

ps_onlyfungi
```

```
## phyloseq-class experiment-level object
## otu_table()   OTU Table:         [ 1889 taxa and 235 samples ]
## sample_data() Sample Data:       [ 235 samples by 37 sample variables ]
## tax_table()   Taxonomy Table:    [ 1889 taxa by 7 taxonomic ranks ]
```

```r
ps_onlyfungi_NC
```

```
## phyloseq-class experiment-level object
## otu_table()   OTU Table:         [ 1851 taxa and 235 samples ]
## sample_data() Sample Data:       [ 235 samples by 37 sample variables ]
## tax_table()   Taxonomy Table:    [ 1851 taxa by 7 taxonomic ranks ]
```

#### Removing leftover plant + arthropod seqs

```r
ps_onlyfungi_NC_OG <- subset_taxa(ps_onlyfungi_NC, Kingdom ==
    "k__Fungi")

ps_onlyfungi_NC_OG
```

```
## phyloseq-class experiment-level object
## otu_table()   OTU Table:         [ 1850 taxa and 235 samples ]
## sample_data() Sample Data:       [ 235 samples by 37 sample variables ]
## tax_table()   Taxonomy Table:    [ 1850 taxa by 7 taxonomic ranks ]
```

#### Remove postitive and negative controls

```r
ps.nocontrols.pear_nomin_err3_v5 <- subset_samples(ps_onlyfungi_NC_OG,
    SampleType != "Control_Buffer")
ps.nocontrols.pear_nomin_err3_v5 <- subset_samples(ps.nocontrols.pear_nomin_err3_v5,
    SampleType != "Control_EtoH")
ps.nocontrols.pear_nomin_err3_v5 <- subset_samples(ps.nocontrols.pear_nomin_err3_v5,
    SampleType != "Control_H20")
```

```r
ps.nocontrols.pear_nomin_err3_v5 <- subset_samples(ps.nocontrols.pear_nomin_err3_v5,
    SampleType != "Control_Kit")
ps.nocontrols.pear_nomin_err3_v5 <- subset_samples(ps.nocontrols.pear_nomin_err3_v5,
    SampleType != "Control_ZymoMockComm")


# remove samples with 0 reads, 4 samples
ps.nocontrols.pear_nomin_err3.v5_nz <- prune_samples(sample_sums(ps.nocontrols.pear_nomin_err3_v5) >
    0, ps.nocontrols.pear_nomin_err3_v5)
```

##### Define standard error function

```r
se <- function(x) sqrt(var(x)/length(x))
```

##### Subset / rarefy samples and filter dataset for bulk sample comparisons

```r
ps.nocontrols.pear_nomin_err3_v5_rare10000 <- rarefy_even_depth(ps.nocontrols.pear_nomin_err3.v5_nz,
    10000, replace = FALSE, rngseed = 5311)
```

```
## `set.seed(5311)` was used to initialize repeatable random subsampling.

## Please record this for your records so others can reproduce.

## Try `set.seed(5311); .Random.seed` for the full vector

## ...

## 31 samples removedbecause they contained fewer reads than `sample.size`.

## Up to first five removed samples are:

## 102.assembled.fastq.gz114.assembled.fastq.gz117.assembled.fastq.gz119.assembled.fastq.gz125.assemble

## ...

## 127OTUs were removed because they are no longer
## present in any sample after random subsampling

## ...
```

```r
# 31 samples removed

# Subset to only bulk samples (LOC)
ps.nocontrols.pear_nomin_err3_rare10000.LOC <- subset_samples(ps.nocontrols.pear_nomin_err3_v5_rare1000
    Project != "COR")

# Remove cores from outside seagrass beds (NEG)
ps.nocontrols.pear_nomin_err3_rare10000.LOC.NoNEG <- subset_samples(ps.nocontrols.pear_nomin_err3_rare1
    Project != "NEG")

# Remove any endophyte samples (no endo)
ps.nocontrols.pear_nomin_err3_rare10000.LOC.NoNEG.noendo <- subset_samples(ps.nocontrols.pear_nomin_err
    SampleSubSubType != "Endophyte")

# Remove any samples from the second location (no GP)
```

```r
ps.nocontrols.pear_nomin_err3_rare10000.LOC.NoNEG.noendo.noGP <- subset_samples(ps.nocontrols.pear_nomi
    Location != "GaffneyPoint")

# how many samples
summary(sample_data(ps.nocontrols.pear_nomin_err3_rare10000.LOC.NoNEG.noendo.noGP)$SampleType)
```

```
##     Leaf  Rhizome     Root Sediment
##       13        7       14       15
```

```r
# Relative abundance
ps.nocontrols.pear_nomin_err3_rare10000.LOC.NoNEG.RA.noendo.noGP <- transform_sample_counts(ps.nocontrol
    function(x) x/sum(x))
```

##### General Leaf, Root, Rhizome, Sediment plot with std err bars

##### Order level

```r
# Order Level
AvgRA_g = tax_glom(ps.nocontrols.pear_nomin_err3_rare10000.LOC.NoNEG.RA.noendo.noGP,
    taxrank = "Order", NArm = FALSE)

AvgRA99_g = filter_taxa(AvgRA_g, function(x) var(x) > 0.01, TRUE)

df_g <- psmelt(AvgRA99_g)

grouped_g <- group_by(df_g, SampleType, SampleSubType, SampleSpecificType,
    Order, Class, Phylum, Kingdom)
avgs_g <- summarise(grouped_g, mean = 100 * mean(Abundance),
    sd = 100 * sd(Abundance), se = 100 * se(Abundance))


avgs_g %<>% mutate(Order2 = fct_explicit_na(Order, na_level = "Unclassified Fungi"))

avgs_g %<>% mutate(Phy2 = fct_explicit_na(Phylum, na_level = "Unclassified Fungi"))

avgs_g$Order2 <- factor(avgs_g$Order2, levels = c("o__Capnodiales",
    "o__Coniochaetales", "o__Dothideales", "o__Eurotiales", "o__Glomerellales",
    "o__Helotiales", "o__Hypocreales", "o__Microascales", "o__Microbotryomycetes_ord_Incertae_sedis",
    "o__Ophiostomatales", "o__Pleosporales", "o__Saccharomycetales",
    "o__Thelebolales", "o__Agaricales", "o__Cystobasidiales",
    "o__Cystofilobasidiales", "o__Filobasidiales", "o__Hymenochaetales",
    "o__Malasseziales", "o__Sporidiobolales", "o__Tremellales",
    "o__Wallemiales", "o__Lobulomycetales", "o__Entorrhizales",
    "Unclassified Fungi"))

avgs_g$SampleType <- factor(avgs_g$SampleType, levels = c("Leaf",
    "Root", "Rhizome", "Sediment"))

p = ggplot(avgs_g, aes(x = Order2, y = (mean), fill = Phy2)) +
    geom_bar(stat = "identity", position = position_dodge()) +
    geom_errorbar(aes(ymin = (mean - se), ymax = (mean + se)),
        width = 0.4, position = position_dodge(0.9))
p = p + facet_grid(~SampleType) + theme(axis.text.x = element_text(angle = -70,
```

```r
    hjust = 0, vjust = 0.5)) + ylab("Mean Relative Abundance") +
    theme(text = element_text(size = 28)) + scale_x_discrete(labels = c(o__Capnodiales = "Capnodiales",
    o__Coniochaetales = "Coniochaetales", o__Eurotiales = "Eurotiales",
    o__Glomerellales = "Glomerellales", o__Helotiales = "Helotiales",
    o__Hypocreales = "Hypocreales", o__Microascales = "Microascales",
    o__Ophiostomatales = "Ophiostomatales", o__Pleosporales = "Pleosporales",
    o__Saccharomycetales = "Saccharomycetales", o__Thelebolales = "Thelebolales",
    o__Agaricales = "Agaricales", o__Cystobasidiales = "Cystobasidiales",
    o__Cystofilobasidiales = "Cystofilobasidiales", o__Filobasidiales = "Filobasidiales",
    o__Hymenochaetales = "Hymenochaetales", o__Malasseziales = "Malasseziales",
    o__Sporidiobolales = "Sporidiobolales", o__Lobulomycetales = "Lobulomycetales",
    `Unclassified Fungi` = "Unclassified Fungi", o__Wallemiales = "Wallemiales",
    o__Tremellales = "Tremellales", o__Microbotryomycetes_ord_Incertae_sedis = "Microbotryomycetes ord.
    o__Entorrhizales = "Entorrhizales", o__Dothideales = "Dothideales"))
p + scale_fill_manual(values = c("#2e919a", "#fdc34a", "#3F5CAA",
    "grey50"), labels = c("Ascomycota", "Basidiomycota", "Chytridiomycota",
    "Unclassified Fungi")) + xlab("Order") + guides(fill = guide_legend(title = "Phylum"))
```

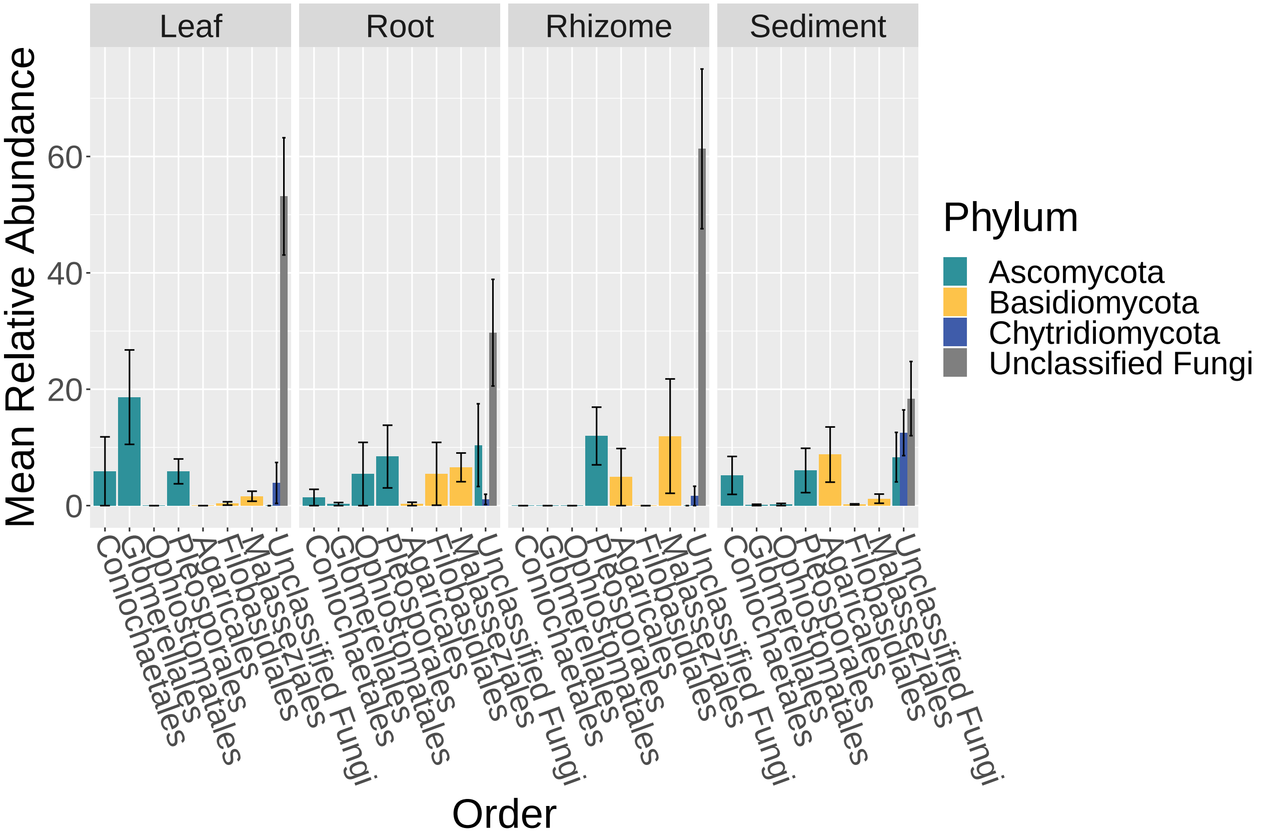


```r
write.csv(avgs_g, "ps.nocontrols.pear_nomin_err3_rare10000.LOC.NoNEG.RA.noendo.noGP_Avgs_var01.csv")
```

##### Stats to investigate which orders vary between bulk sample types

```r
# start
chisq = NULL
pvals = NULL
listofcats_sig = NULL
```

```r
DataSet = df_g

# remove NAs
DataSet %<>% mutate(Order2 = fct_explicit_na(Order, na_level = "Unclassified"))
taxa = unique(DataSet$Order2)

for (cat in taxa) {
    new_df <- subset(DataSet, DataSet$Order2 == cat)
    new_df$ST <- as.factor(new_df$SampleType)
    kw = kruskal.test(Abundance ~ ST, data = new_df)
    chisq = c(kw$statistic, chisq)
    pvals = c(kw$p.value, pvals)

    if (kw$p.value <= 0.05) {
        listofcats_sig = c(cat, listofcats_sig)

    }
}

pvals.bonf = p.adjust(pvals, method = "bonferroni")
df_taxa = data.frame(rev(taxa), chisq, pvals, pvals.bonf)
write.table(df_taxa, "SampleType_rare10000_Mean_KW_01var.txt",
    sep = "\t")


pvals.dunn = NULL
pvals.dunn.bonf = NULL
Zsc.dunn = NULL
comparison.dunn = NULL
cats.dunn = NULL

for (cat in listofcats_sig) {
    new_df <- subset(DataSet, DataSet$Order2 == cat)
    new_df$ST <- as.factor(new_df$SampleType)
    dT = dunnTest(Abundance ~ ST, data = new_df, method = "bonferroni")

    for (i in 1:length(dT$res$Comparison)) {
        print(cat)

        pvals.dunn = c(dT$res$P.unadj[i], pvals.dunn)
        pvals.dunn.bonf = c(dT$res$P.adj[i], pvals.dunn.bonf)
        Zsc.dunn = c(dT$res$Z[i], Zsc.dunn)
        comparison.dunn = c(as.character(dT$res$Comparison)[i],
            comparison.dunn)
        cats.dunn = c(cat, cats.dunn)

    }

}


df_taxa = data.frame(cats.dunn, comparison.dunn, Zsc.dunn, pvals.dunn,
    pvals.dunn.bonf)
```

```
write.table(df_taxa, "SampleType_rare10000_Mean_order_KW_01var_Dunn.txt",
    sep = "\t")
```

##### Ordination of bulk sample types

```
pp_pcoa <- ordinate(physeq = ps.nocontrols.pear_nomin_err3_rare10000.LOC.NoNEG.noendo.noGP,
    method = "PCoA", distance = "bray")

p = plot_ordination(physeq = ps.nocontrols.pear_nomin_err3_rare10000.LOC.NoNEG.noendo.noGP,
    ordination = pp_pcoa, shape = "SampleType", color = "SampleType")
p = p + geom_point(size = 6) + theme(text = element_text(size = 28)) +
    labs(color = "Sample Type", shape = "Sample Type")  #+ scale_color_manual(values=c('#39c44a','#3F5C
p + xlab("PCoA 1 (10.4% of variation)") + ylab("PCoA 2 (9.1% of variation)")
```

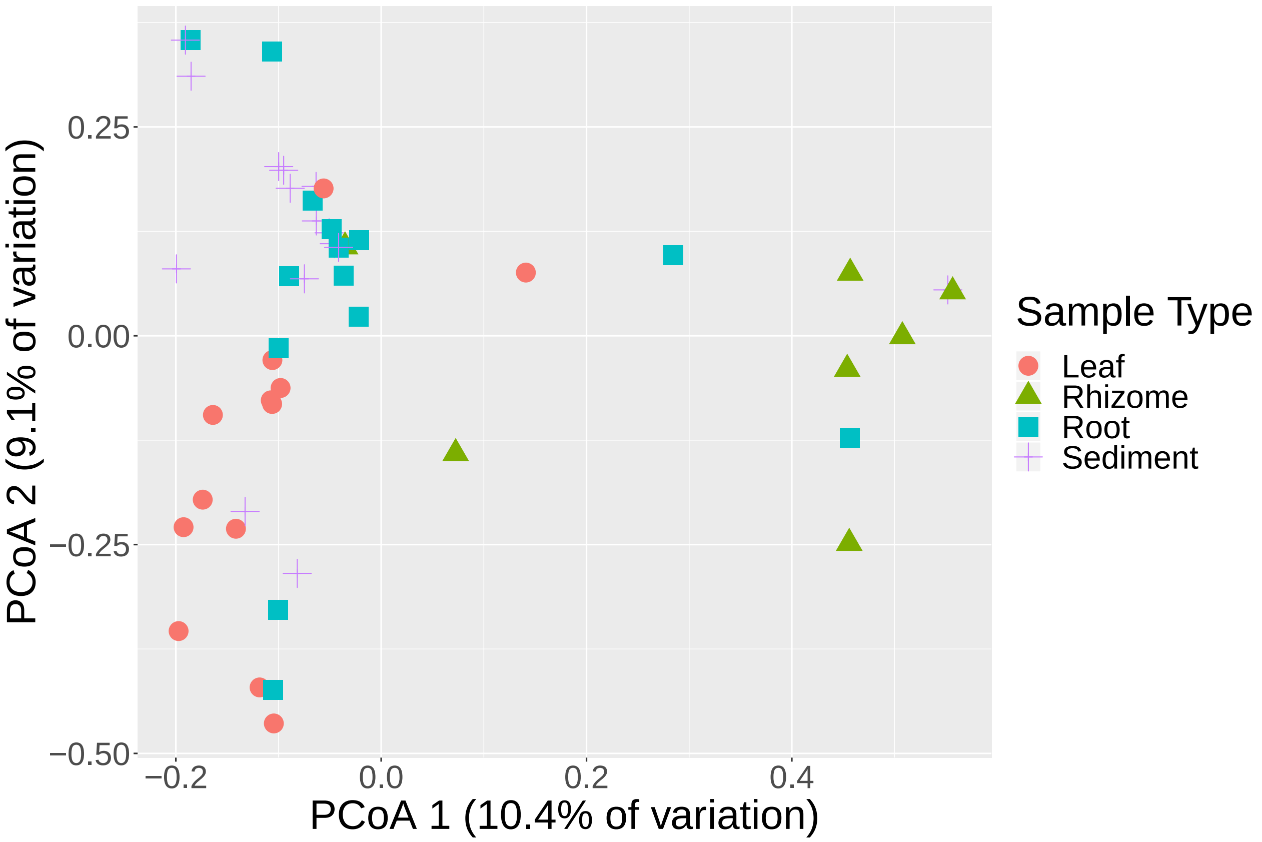


#### Ordination Stats

```
listofcats = colnames(as(sample_data(ps.nocontrols.pear_nomin_err3_rare10000.LOC.NoNEG.noendo.noGP),
    "data.frame"))
listofcats = listofcats[listofcats != "Notes"]
listofcats = listofcats[listofcats != "Location"]
listofcats = listofcats[listofcats != "Project"]
listofcats = listofcats[listofcats != "CNRatio_PlantTissue"]
listofcats = listofcats[listofcats != "IsControl"]
listofcats = listofcats[listofcats != "SampleSubType"]
listofcats = listofcats[listofcats != "WeightTissue..g."]
listofcats = listofcats[listofcats != "DNA.Concentration..ng.ul."]
```

```
listofcats = listofcats[listofcats != "WeightORVolumeExtracted..g.or.ul."]
listofcats = listofcats[listofcats != "Lattitude"]
listofcats = listofcats[listofcats != "Longitude"]
listofcats = listofcats[listofcats != "Description"]
listofcats = listofcats[listofcats != "ExtractionKitType"]
listofcats = listofcats[listofcats != "Salinity_H20"]
listofcats = listofcats[listofcats != "pH_H20"]
listofcats = listofcats[listofcats != "Nitrate_H20_mgperL"]
listofcats = listofcats[listofcats != "Nitrite_H20_ppb"]
listofcats = listofcats[listofcats != "Phosphorous_H20"]
listofcats = listofcats[listofcats != "Ammonia_H20_mgperL"]
listofcats = listofcats[listofcats != "SampleID"]
listofcats = listofcats[listofcats != "OriginalSampleID"]
listofcats = listofcats[listofcats != "RealSampleID"]
listofcats = listofcats[listofcats != "SampleNumber"]
listofcats = listofcats[listofcats != "DateWeighed"]
listofcats = listofcats[listofcats != "ZymoShipLocation"]
listofcats = listofcats[listofcats != "ZymoSampleID"]
listofcats = listofcats[listofcats != "ZymoWell"]
listofcats = listofcats[listofcats != "SampleSpecificType"]
listofcats = listofcats[listofcats != "SampleSubSubType"]
listofcats = listofcats[listofcats != "CollectionDate"]
listofcats = listofcats[listofcats != "ExtractionDate"]

All.distBC = phyloseq::distance(ps.nocontrols.pear_nomin_err3_rare10000.LOC.NoNEG.noendo.noGP,
    method = "bray", type = "samples")
DataSet = ps.nocontrols.pear_nomin_err3_rare10000.LOC.NoNEG.noendo.noGP

pvals = NULL
Fmods = NULL
Rs = NULL
listofcats_sig = NULL

for (cat in listofcats) {
    form <- as.formula(paste("All.distBC", cat, sep = "~"))
    results = adonis(form, as(sample_data(DataSet), "data.frame"),
        permutations = 9999)
    pvals = c(results$aov.tab$"Pr(>F)"[1], pvals)
    Fmods = c(results$aov.tab$F.Model[1], Fmods)
    Rs = c(results$aov.tab$R2[1], Rs)


    if (results$aov.tab$"Pr(>F)"[1] <= 0.05) {

        listofcats_sig = c(cat, listofcats_sig)
    }

}

pvals.bonf = p.adjust(pvals, method = "bonferroni")
df = data.frame(rev(listofcats), Fmods, Rs, pvals, pvals.bonf)
write.table(df, "ps.nocontrols.pear_nomin_err3_rare10000.LOC.NoNEG_noendo_permanova_BC.txt",
    sep = "\t")
```

```r
# Categories with significant PERMANOVA results (before
# p-value correction)
listofcats_sig
# [1] 'CoreNumber' 'TimePoint' 'SampleType'

## Sample Type Pair-wise permanovas - if was to redo today
## would use the function in EcoUtils package instead (didn't
## exist at the time)
cat.sub = unique(as(sample_data(DataSet), "data.frame")$SampleType)

pvals.sub = NULL
Fmods.sub = NULL
Rs.sub = NULL

# pw_combos ends up giving you the factor #
pw_combos = combn(cat.sub, 2)
pw_1 = NULL
pw_2 = NULL

i = 1
for (i in (1:((length(pw_combos) + 1)/2))) {
    pw_1 = c(pw_combos[, i][1], pw_1)
    pw_2 = c(pw_combos[, i][2], pw_2)
    sub.biom1 <- subset_samples(DataSet, SampleType == pw_combos[,
        i][1])
    sub.biom2 <- subset_samples(DataSet, SampleType == pw_combos[,
        i][2])
    pw.biom = merge_phyloseq(sub.biom1, sub.biom2)

    i = i + 1
    sub.distBC = phyloseq::distance(pw.biom, method = "bray",
        type = "samples")

    result.sub = adonis(sub.distBC ~ SampleType, as(sample_data(pw.biom),
        "data.frame"), permutations = 9999)

    pvals.sub = c(result.sub$aov.tab$"Pr(>F)"[1], pvals.sub)
    Fmods.sub = c(result.sub$aov.tab$F.Model[1], Fmods.sub)
    Rs.sub = c(result.sub$aov.tab$R2[1], Rs.sub)
}

pvals.sub.bonf = p.adjust(pvals.sub, method = "bonferroni")
sub.df = data.frame(pw_1, pw_2, Fmods.sub, Rs.sub, pvals.sub,
    pvals.sub.bonf)

write.table(sub.df, "SampleType_ps.nocontrols.pear_nomin_err3_rare10000.LOC.NoNEG_noendo_pw_permanova_B
    sep = "\t")


# betadisper on above Dataset
pvals = NULL
Fmods = NULL
Rs = NULL
```

```
listofcats_sig = NULL

for (cat in listofcats) {
    new_cat <- as.character(cat)
    disp_dist <- betadisper(All.distBC, as(sample_data(DataSet),
        "data.frame")[, cat])
    results = permutest(disp_dist, permutations = 9999)
    pvals = c(results$tab$"Pr(>F)"[1], pvals)
    Fmods = c(results$tab$F[1], Fmods)

    if (results$tab$"Pr(>F)"[1] <= 0.05) {

        listofcats_sig = c(cat, listofcats_sig)
    }

}

pvals.bonf = p.adjust(pvals, method = "bonferroni")
df = data.frame(rev(listofcats), Fmods, pvals, pvals.bonf)
write.table(df, "ps.nocontrols.pear_nomin_err3_rare10000.LOC.NoNEG.noendo.noGP_betadisper_BC.txt",
    sep = "\t")

listofcats_sig
#'RandomizedExtractionGroup' 'SampleType'

disp_dist <- betadisper(All.distBC, as(sample_data(DataSet),
    "data.frame")$SampleType)
TukeyHSD(disp_dist)

disp_dist <- betadisper(All.distBC, as(sample_data(DataSet),
    "data.frame")$RandomizedExtractionGroup)
TukeyHSD(disp_dist)
```

##### Alpha Diversity of bulk samples

```
p = plot_richness(ps.nocontrols.pear_nomin_err3_rare10000.LOC.NoNEG.noendo.noGP,
    measures = c("Observed", "Shannon"), x = "SampleType", color = "SampleType")
p = p + theme(text = element_text(size = 24)) + geom_boxplot() +
    geom_jitter() + theme(axis.text.x = element_text(angle = -70,
    hjust = 0, vjust = 0.5))
p + scale_colour_manual(values = c("#2e919a", "#fdc34a", "#3F5CAA",
    "grey50")) + xlab("Sample Type") + guides(color = guide_legend(title = "Sample Type"))
```

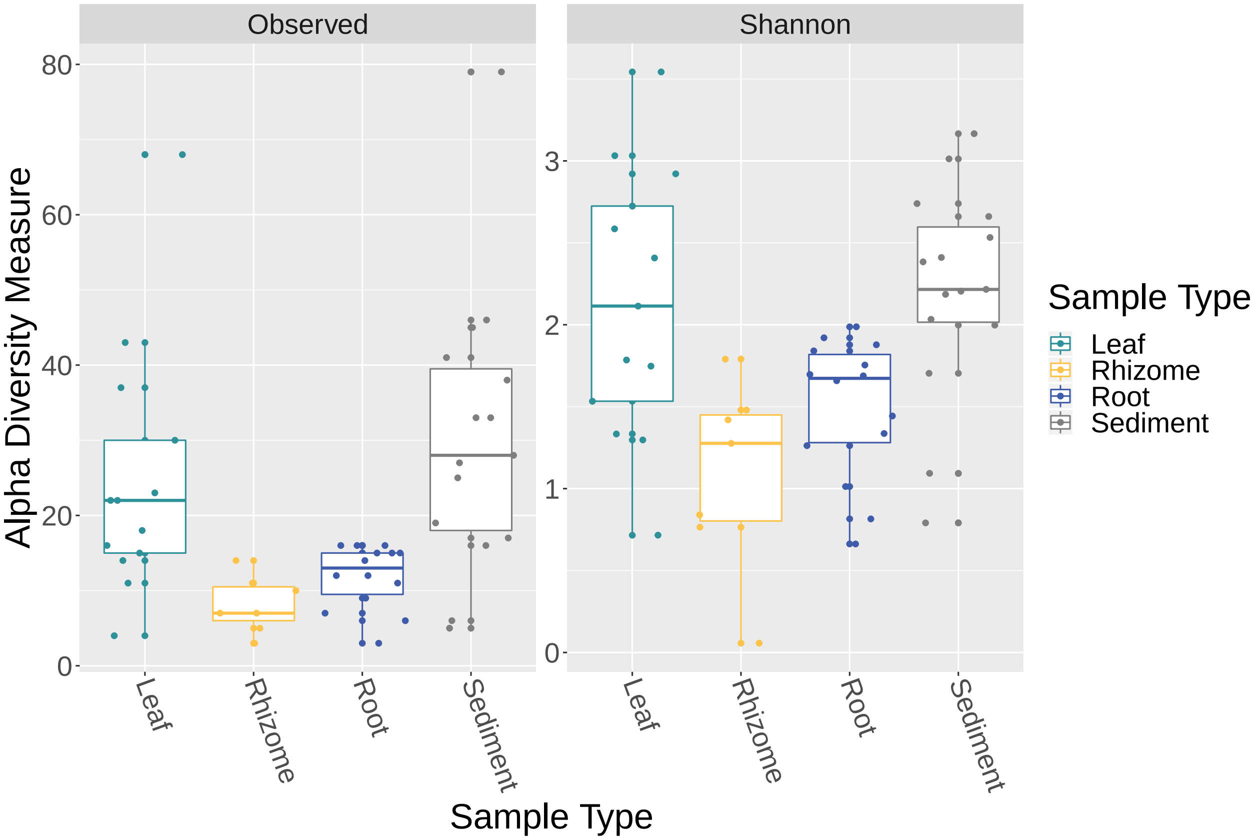


##### Stats for alpha diversity across bulk samples

```
LOC_Alpha = estimate_richness(ps.nocontrols.pear_nomin_err3_rare10000.LOC.NoNEG.noendo.noGP,
    measures = c("Observed", "Shannon"))

LOC_Alpha_2 <- cbind(LOC_Alpha, sample_data(ps.nocontrols.pear_nomin_err3_rare10000.LOC.NoNEG.noendo.no

# Sample Type
LOC_Alpha_2$ST <- as.factor(LOC_Alpha_2$SampleType)

# KW tests on alpha diversity across sample types
kruskal_test(Observed ~ ST, distribution = approximate(nresample = 9999),
    data = LOC_Alpha_2)
```

```
##
##  Approximative Kruskal-Wallis Test
##
## data:  Observed by ST (Leaf, Rhizome, Root, Sediment)
## chi-squared = 20.632, p-value = 1e-04
```

```
kruskal_test(Shannon ~ ST, distribution = approximate(nresample = 9999),
    data = LOC_Alpha_2)
```

```
##
##  Approximative Kruskal-Wallis Test
##
```

```
## data:  Shannon by ST (Leaf, Rhizome, Root, Sediment)
## chi-squared = 16.101, p-value = 0.0007001
```

```
# Post-hoc tests
dunnTest(Observed ~ ST, data = LOC_Alpha_2, method = "bonferroni")
```

```
## Dunn (1964) Kruskal-Wallis multiple comparison

##   p-values adjusted with the Bonferroni method.

##           Comparison          Z      P.unadj        P.adj
## 1     Leaf - Rhizome  3.0394389 0.0023701927 0.0142211564
## 2        Leaf - Root  2.3415425 0.0192042391 0.1152254346
## 3     Rhizome - Root -1.1298783 0.2585275092 1.0000000000
## 4    Leaf - Sediment -0.8397157 0.4010678164 1.0000000000
## 5 Rhizome - Sediment -3.8080900 0.0001400443 0.0008402658
## 6    Root - Sediment -3.2831971 0.0010263688 0.0061582129
```

```
dunnTest(Shannon ~ ST, data = LOC_Alpha_2, method = "bonferroni")
```

```
## Dunn (1964) Kruskal-Wallis multiple comparison
##   p-values adjusted with the Bonferroni method.

##           Comparison          Z      P.unadj       P.adj
## 1     Leaf - Rhizome  2.7510910 0.0059397148 0.035638289
## 2        Leaf - Root  2.1544749 0.0312029459 0.187217675
## 3     Rhizome - Root -0.9935066 0.3204631285 1.000000000
## 4    Leaf - Sediment -0.6156369 0.5381342165 1.000000000
## 5 Rhizome - Sediment -3.3272681 0.0008770195 0.005262117
## 6    Root - Sediment -2.8608140 0.0042255492 0.025353295
```

#### Subset / rarefy samples for leaf length comparisons

```
# Rarefied data
ps.nocontrols.pear_nomin_err3_v5_rare5000 <- rarefy_even_depth(ps.nocontrols.pear_nomin_err3.v5_nz,
    5000, replace = FALSE, rngseed = 5311)
```

```
## `set.seed(5311)` was used to initialize repeatable random subsampling.

## Please record this for your records so others can reproduce.

## Try `set.seed(5311); .Random.seed` for the full vector

## ...

## 22 samples removedbecause they contained fewer reads than `sample.size`.

## Up to first five removed samples are:

## 114.assembled.fastq.gz117.assembled.fastq.gz119.assembled.fastq.gz125.assembled.fastq.gz147.assemble

## ...

## 162OTUs were removed because they are no longer
## present in any sample after random subsampling

## ...
```

```
# Only intra-plant samples
ps.nocontrols.pear_nomin_err3.COR.rare5000 <- subset_samples(ps.nocontrols.pear_nomin_err3_v5_rare5000,
    Project != "LOC")
```

```r
# No negative control cores (NEG)
ps.nocontrols.pear_nomin_err3.COR.rare5000.NoNEG <- subset_samples(ps.nocontrols.pear_nomin_err3.COR.ra
    Project != "NEG")

# Only leaf samples
ps.nocontrols.pear_nomin_err3.COR.rare5000.NoNEG.LEAF <- subset_samples(ps.nocontrols.pear_nomin_err3.C
    SampleType == "Leaf")

# how many samples
summary(sample_data(ps.nocontrols.pear_nomin_err3.COR.rare5000.NoNEG.LEAF)$SampleSubType)
```

```
## Endophyte  Epiphyte
##        25        25
```

```r
# Relative abundance
ps.nocontrols.pear_nomin_err3.COR.rare5000.NoNEG.LEAF.RA <- transform_sample_counts(ps.nocontrols.pear_
    function(x) x/sum(x))


# Only root samples
ps.nocontrols.pear_nomin_err3.COR.NoNEG.rare5000.ROOT <- subset_samples(ps.nocontrols.pear_nomin_err3.C
    SampleType == "Root")

# how many samples
summary(sample_data(ps.nocontrols.pear_nomin_err3.COR.NoNEG.rare5000.ROOT)$SampleSubType)
```

```
## Endophyte  Epiphyte
##        14        21
```

##### Ordination of leaf epiphytes vs. endophytes

```r
loc_pcoa <- ordinate(physeq = ps.nocontrols.pear_nomin_err3.COR.rare5000.NoNEG.LEAF,
    method = "PCoA", distance = "bray")

## FIGURE for EPI v ENDO
p = plot_ordination(physeq = ps.nocontrols.pear_nomin_err3.COR.rare5000.NoNEG.LEAF,
    ordination = loc_pcoa, shape = "SampleSubType", color = "SampleSubType")
p = p + geom_point(size = 6) + theme(text = element_text(size = 28)) +
    labs(color = "Sample Type", shape = "Sample Type") + scale_color_manual(values = c("#39c44a",
    "#3F5CAA"))
p + xlab("PCoA 1 (28.1% of variation)") + ylab("PCoA 2 (14.5% of variation)")
```

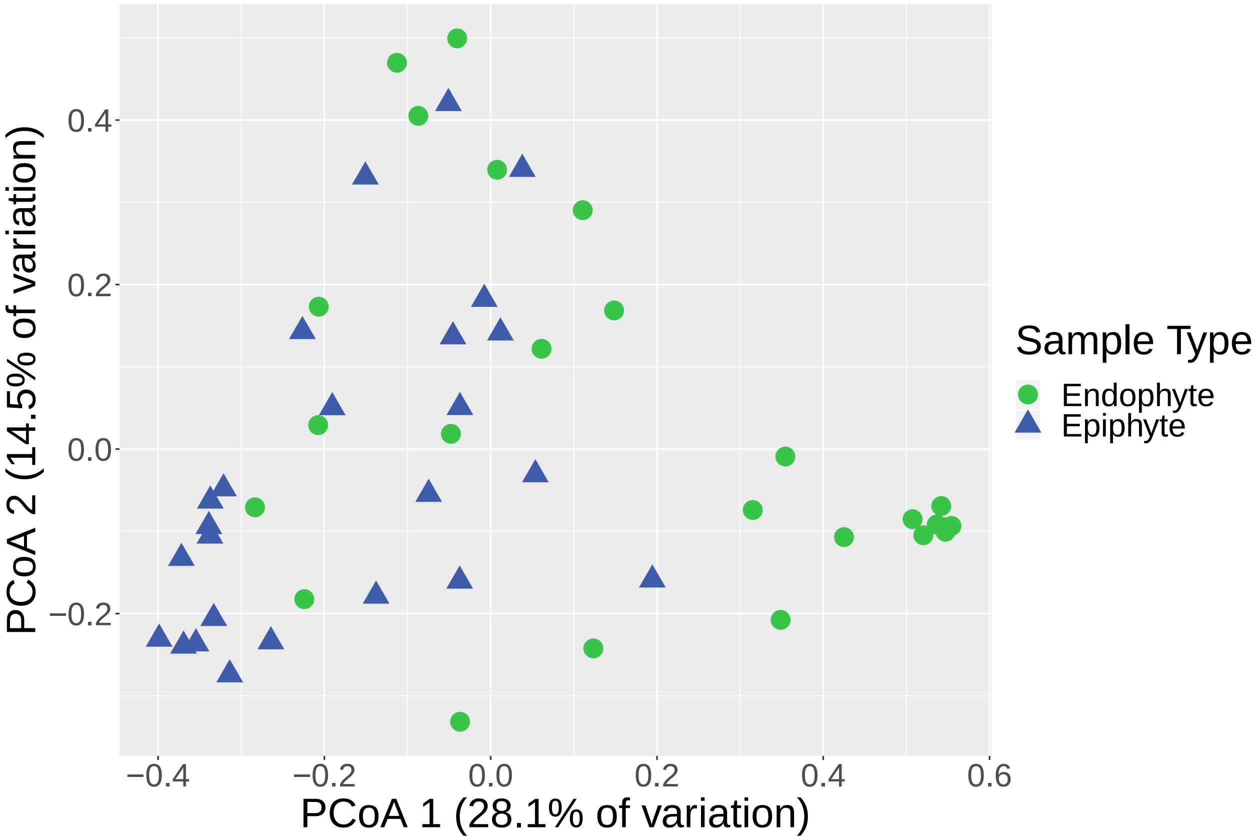


##### Stats on leaf epi vs. endo ordination

```
listofcats = colnames(as(sample_data(ps.nocontrols.pear_nomin_err3.COR.rare5000.NoNEG.LEAF),
    "data.frame"))
listofcats = listofcats[listofcats != "Notes"]
listofcats = listofcats[listofcats != "Location"]
listofcats = listofcats[listofcats != "Project"]
listofcats = listofcats[listofcats != "CNRatio_PlantTissue"]
listofcats = listofcats[listofcats != "IsControl"]
listofcats = listofcats[listofcats != "WeightTissue..g."]
listofcats = listofcats[listofcats != "DNA.Concentration..ng.ul."]
listofcats = listofcats[listofcats != "WeightORVolumeExtracted..g.or.ul."]
listofcats = listofcats[listofcats != "Lattitude"]
listofcats = listofcats[listofcats != "Longitude"]
listofcats = listofcats[listofcats != "Description"]
listofcats = listofcats[listofcats != "ExtractionKitType"]
listofcats = listofcats[listofcats != "Salinity_H20"]
listofcats = listofcats[listofcats != "pH_H20"]
listofcats = listofcats[listofcats != "Nitrate_H20_mgperL"]
listofcats = listofcats[listofcats != "Nitrite_H20_ppb"]
listofcats = listofcats[listofcats != "Phosphorous_H20"]
listofcats = listofcats[listofcats != "Ammonia_H20_mgperL"]
listofcats = listofcats[listofcats != "SampleID"]
listofcats = listofcats[listofcats != "OriginalSampleID"]
listofcats = listofcats[listofcats != "RealSampleID"]
```

```r
listofcats = listofcats[listofcats != "SampleNumber"]
listofcats = listofcats[listofcats != "DateWeighed"]
listofcats = listofcats[listofcats != "ZymoShipLocation"]
listofcats = listofcats[listofcats != "ZymoSampleID"]
listofcats = listofcats[listofcats != "ZymoWell"]
listofcats = listofcats[listofcats != "SampleType"]
listofcats = listofcats[listofcats != "SampleSubSubType"]
listofcats = listofcats[listofcats != "CollectionDate"]
listofcats = listofcats[listofcats != "ExtractionDate"]
listofcats = listofcats[listofcats != "TimePoint"]

All.distBC = phyloseq::distance(ps.nocontrols.pear_nomin_err3.COR.rare5000.NoNEG.LEAF,
    method = "bray", type = "samples")
DataSet = ps.nocontrols.pear_nomin_err3.COR.rare5000.NoNEG.LEAF

pvals = NULL
Fmods = NULL
Rs = NULL
listofcats_sig = NULL

for (cat in listofcats) {
    form <- as.formula(paste("All.distBC", cat, sep = "~"))
    results = adonis(form, as(sample_data(DataSet), "data.frame"),
        permutations = 9999)
    pvals = c(results$aov.tab$"Pr(>F)"[1], pvals)
    Fmods = c(results$aov.tab$F.Model[1], Fmods)
    Rs = c(results$aov.tab$R2[1], Rs)


    if (results$aov.tab$"Pr(>F)"[1] <= 0.05) {

        listofcats_sig = c(cat, listofcats_sig)
    }

}

pvals.bonf = p.adjust(pvals, method = "bonferroni")
df = data.frame(rev(listofcats), Fmods, Rs, pvals, pvals.bonf)
write.table(df, "ps.nocontrols.pear_nomin_err3.COR.rare5000.NoNEG.LEAF_permanova_BC.txt",
    sep = "\t")

# samplespecifictype
cat.sub = unique(as(sample_data(DataSet), "data.frame")$SampleSpecificType)

pvals.sub = NULL
Fmods.sub = NULL
Rs.sub = NULL

# pw_combos ends up giving you the factor #
pw_combos = combn(cat.sub, 2)
pw_1 = NULL
pw_2 = NULL
```

```r
i = 1
for (i in (1:((length(pw_combos) + 1)/2))) {
    pw_1 = c(pw_combos[, i][1], pw_1)
    pw_2 = c(pw_combos[, i][2], pw_2)
    sub.biom1 <- subset_samples(DataSet, SampleSpecificType ==
        pw_combos[, i][1])
    sub.biom2 <- subset_samples(DataSet, SampleSpecificType ==
        pw_combos[, i][2])
    pw.biom = merge_phyloseq(sub.biom1, sub.biom2)

    i = i + 1
    sub.distBC = phyloseq::distance(pw.biom, method = "bray",
        type = "samples")

    result.sub = adonis(sub.distBC ~ SampleSpecificType, as(sample_data(pw.biom),
        "data.frame"), permutations = 9999)

    pvals.sub = c(result.sub$aov.tab$"Pr(>F)"[1], pvals.sub)
    Fmods.sub = c(result.sub$aov.tab$F.Model[1], Fmods.sub)
    Rs.sub = c(result.sub$aov.tab$R2[1], Rs.sub)
}

pvals.sub.bonf = p.adjust(pvals.sub, method = "bonferroni")
sub.df = data.frame(pw_1, pw_2, Fmods.sub, Rs.sub, pvals.sub,
    pvals.sub.bonf)

write.table(sub.df, "SampleSpecificType_ps.nocontrols.pear_nomin_err3.COR.rare5000.NoNEG.LEAF_pw_perman
    sep = "\t")


# betadisper on above Dataset
pvals = NULL
Fmods = NULL
Rs = NULL
listofcats_sig = NULL

for (cat in listofcats) {
    new_cat <- as.character(cat)
    disp_dist <- betadisper(All.distBC, as(sample_data(DataSet),
        "data.frame")[, cat])
    results = permutest(disp_dist, permutations = 9999)
    pvals = c(results$tab$"Pr(>F)"[1], pvals)
    Fmods = c(results$tab$F[1], Fmods)

    if (results$tab$"Pr(>F)"[1] <= 0.05) {

        listofcats_sig = c(cat, listofcats_sig)
    }

}

pvals.bonf = p.adjust(pvals, method = "bonferroni")
df = data.frame(rev(listofcats), Fmods, pvals, pvals.bonf)
```

```r
write.table(df, "ps.nocontrols.pear_nomin_err3.COR.rare5000.NoNEG.LEAF_betadisper_BC.txt",
    sep = "\t")

listofcats_sig


disp_dist <- betadisper(All.distBC, as(sample_data(DataSet),
    "data.frame")$SampleSpecificType)
TukeyHSD(disp_dist)
```

##### Investigating orders that vary across leaf length

```r
AvgRA_leaf = tax_glom(ps.nocontrols.pear_nomin_err3.COR.rare5000.NoNEG.LEAF.RA,
    taxrank = "Order", NArm = FALSE)

AvgRA99_leaf = filter_taxa(AvgRA_leaf, function(x) var(x) > 0.001,
    TRUE)

df_leaf <- psmelt(AvgRA99_leaf)

grouped_leaf <- group_by(df_leaf, SampleType, SampleSpecificType,
    OTU, Order, Class, Phylum)
avgs_leaf <- summarise(grouped_leaf, mean = 100 * mean(Abundance),
    sd = 100 * sd(Abundance), se = 100 * se(Abundance))

avgs_leaf <- read_csv("ps.nocontrols.pear_nomin_err3.COR.RA.NoNEG.rare5000.LEAF_avgs_var001.csv")
```

```
## Parsed with column specification:
## cols(
##   X1 = col_double(),
##   SampleType = col_character(),
##   SampleSpecificType = col_character(),
##   OTU = col_character(),
##   Order = col_character(),
##   Class = col_character(),
##   Phylum = col_character(),
##   mean = col_double(),
##   sd = col_double(),
##   se = col_double(),
##   Phyla2 = col_character(),
##   Order2 = col_character()
## )
```

```r
avgs_leaf %<>% mutate(Phyla2 = fct_explicit_na(Phylum, na_level = "Unclassified Fungi"))
avgs_leaf %<>% mutate(Order2 = fct_explicit_na(Order, na_level = "Unclassified Fungi"))
facet_names <- c(`0_to_5_inches` = "0-5 inches", `10_to_15_inches` = "10-15 inches",
    `15_to_20_inches` = "15-20 inches", `20_to_25_inches` = "20-25 inches",
    `5_to_10_inches` = "5-10 inches")
avgs_leaf$SampleSpecificType <- factor(avgs_leaf$SampleSpecificType,
    levels = c("0_to_5_inches", "5_to_10_inches", "10_to_15_inches",
        "15_to_20_inches", "20_to_25_inches"))
```

```r
avgs_leaf$Order2 <- factor(avgs_leaf$Order2, levels = c("o__Chaetothyriales",
    "o__Glomerellales", "o__Pleosporales", "o__Saccharomycetales",
    "o__Tubeufiales", "o__Malasseziales", "Unclassified Fungi"))

## Fig
p = ggplot(avgs_leaf, aes(x = Order2, y = (mean), fill = Phyla2)) +
    geom_bar(stat = "identity", position = position_dodge()) +
    geom_errorbar(aes(ymin = (mean - se), ymax = (mean + se)),
        width = 0.4, position = position_dodge(0.9))
p = p + facet_grid(~SampleSpecificType, labeller = as_labeller(facet_names)) +
    theme(axis.text.x = element_text(angle = -70, hjust = 0,
        vjust = 0.5)) + ylab("Mean Relative Abundance") + theme(text = element_text(size = 28))
p + xlab("Order") + guides(fill = guide_legend(title = "Phylum")) +
    scale_fill_manual(values = c("#2e919a", "#fdc34a", "grey50"),
        labels = c("Ascomycota", "Basidiomycota", "Unclassified Fungi")) +
    scale_x_discrete(labels = c(o__Chaetothyriales = "Chaetothyriales",
        o__Tubeufiales = "Tubeufiales", o__Glomerellales = "Glomerellales",
        o__Pleosporales = "Pleosporales", o__Saccharomycetales = "Saccharomycetales",
        o__Malasseziales = "Malasseziales", `Unclassified Fungi` = "Unclassified Fungi"))
```

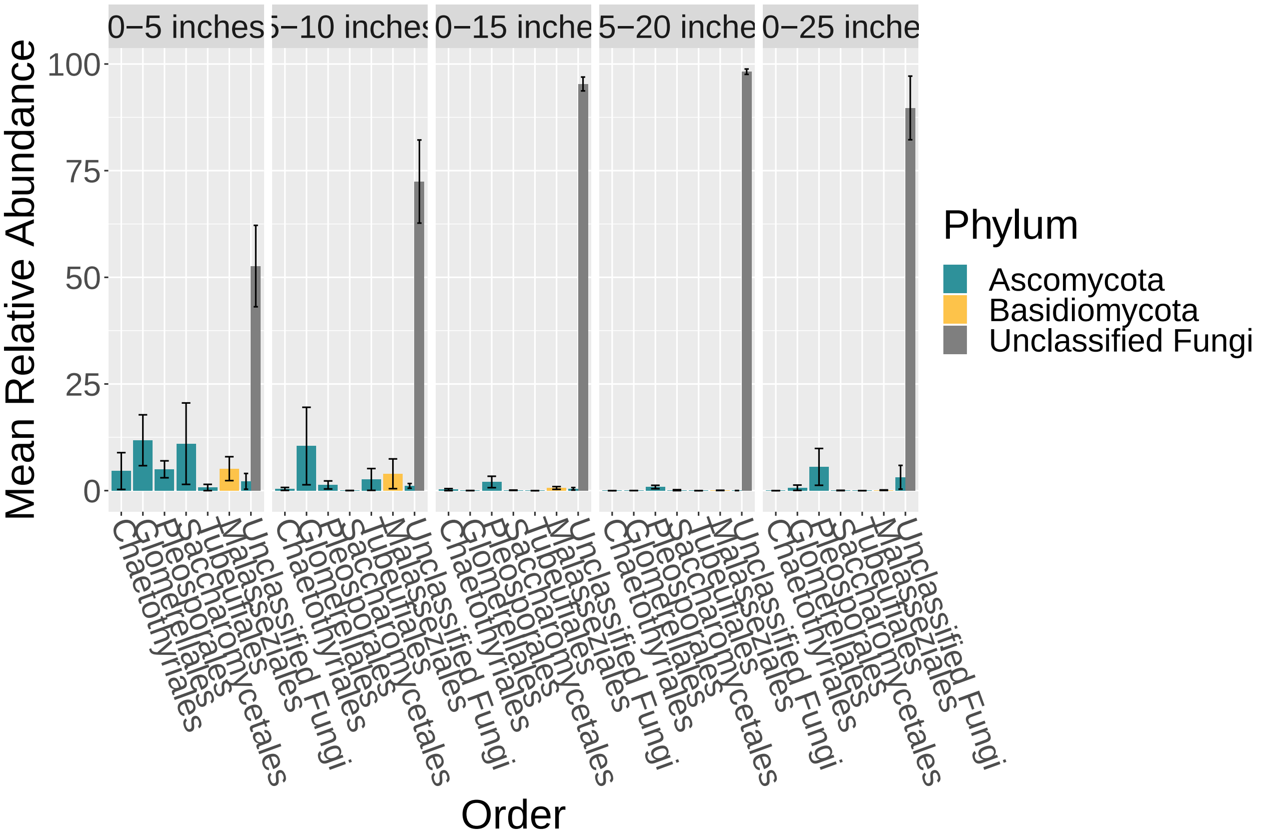


```r
# write.csv(avgs_leaf,
# 'ps.nocontrols.pear_nomin_err3.COR.RA.NoNEG.rare5000.LEAF_avgs_var001.csv')
```

#### Stats on orders that vary across leaf length

```r
# start
chisq = NULL
pvals = NULL
listofcats_sig = NULL
DataSet = df_leaf

# remove NAs
DataSet %<>% mutate(Order2 = fct_explicit_na(Order, na_level = "Unclassified"))
taxa = unique(DataSet$Order2)

for (cat in taxa) {
    new_df <- subset(DataSet, DataSet$Order2 == cat)
    new_df$ST <- as.factor(new_df$SampleSpecificType)
    kw = kruskal.test(Abundance ~ ST, data = new_df)
    chisq = c(kw$statistic, chisq)
    pvals = c(kw$p.value, pvals)

    if (kw$p.value <= 0.05) {
        listofcats_sig = c(cat, listofcats_sig)

    }
}

pvals.bonf = p.adjust(pvals, method = "bonferroni")
df_taxa = data.frame(rev(taxa), chisq, pvals, pvals.bonf)
write.table(df_taxa, "LeafLength_rare5000_Mean_KW_001var.txt",
    sep = "\t")


pvals.dunn = NULL
pvals.dunn.bonf = NULL
Zsc.dunn = NULL
comparison.dunn = NULL
cats.dunn = NULL

for (cat in listofcats_sig) {
    new_df <- subset(DataSet, DataSet$Order2 == cat)
    new_df$ST <- as.factor(new_df$SampleSpecificType)
    dT = dunnTest(Abundance ~ ST, data = new_df, method = "bonferroni")

    for (i in 1:length(dT$res$Comparison)) {
        print(cat)

        pvals.dunn = c(dT$res$P.unadj[i], pvals.dunn)
        pvals.dunn.bonf = c(dT$res$P.adj[i], pvals.dunn.bonf)
        Zsc.dunn = c(dT$res$Z[i], Zsc.dunn)
        comparison.dunn = c(as.character(dT$res$Comparison)[i],
            comparison.dunn)
        cats.dunn = c(cat, cats.dunn)

    }
```

```
}


df_taxa = data.frame(cats.dunn, comparison.dunn, Zsc.dunn, pvals.dunn,
    pvals.dunn.bonf)
write.table(df_taxa, "LeafLength_rare5000_Mean_order_KW_001var_Dunn.txt",
    sep = "\t")
```

##### ASVs that differ across leaf length

```
# ASV by mean
AvgRA99_leaf = filter_taxa(ps.nocontrols.pear_nomin_err3.COR.rare5000.NoNEG.LEAF.RA,
    function(x) mean(x) > 0.02, TRUE)

# Lobulomycetales - SV8, SV11, SV16, SV56 Glomerellales -
# SV10 Aphelidiomycota - SV12 Chytridiomycetes - SV21
# Chytridiomycota - SV28, SV36

# SV8 complex = SV8, SV11, SV16, SV56 SV28 complex = SV28,
# SV36

avgra99_leaf_merge <- merge_taxa(AvgRA99_leaf, c("SV8", "SV11",
    "SV16", "SV56"), "SV8")
avgra99_leaf_merge <- merge_taxa(avgra99_leaf_merge, c("SV28",
    "SV36"), "SV28")

df_leaf_asv <- psmelt(avgra99_leaf_merge)

grouped_leaf <- group_by(df_leaf_asv, SampleType, SampleSpecificType,
    OTU, Order, Class, Phylum)
avgs_leaf <- summarise(grouped_leaf, mean = 100 * mean(Abundance),
    sd = 100 * sd(Abundance), se = 100 * se(Abundance))

avgs_leaf %<>% mutate(Phyla2 = fct_explicit_na(Phylum, na_level = "Unclassified Fungi"))
avgs_leaf %<>% mutate(Order2 = fct_explicit_na(Order, na_level = "Unclassified Fungi"))
facet_names <- c(`0_to_5_inches` = "0-5 inches", `10_to_15_inches` = "10-15 inches",
    `15_to_20_inches` = "15-20 inches", `20_to_25_inches` = "20-25 inches",
    `5_to_10_inches` = "5-10 inches")
avgs_leaf$SampleSpecificType <- factor(avgs_leaf$SampleSpecificType,
    levels = c("0_to_5_inches", "5_to_10_inches", "10_to_15_inches",
        "15_to_20_inches", "20_to_25_inches"))

p = ggplot(avgs_leaf, aes(x = OTU, y = (mean), fill = Phyla2)) +
    geom_bar(stat = "identity", position = position_dodge()) +
    geom_errorbar(aes(ymin = (mean - se), ymax = (mean + se)),
        width = 0.4, position = position_dodge(0.9))
p = p + facet_grid(~SampleSpecificType, labeller = as_labeller(facet_names)) +
    theme(axis.text.x = element_text(angle = -70, hjust = 0,
        vjust = 0.5)) + ylab("Mean Relative Abundance") + theme(text = element_text(size = 28))
p + xlab("ASV") + guides(fill = guide_legend(title = "Phylum")) +
    scale_fill_manual(values = c("#2e919a", "grey50"), labels = c("Ascomycota",
```

```
        "Unclassified Fungi")) + scale_x_discrete(labels = c(SV10 = "SV10",
    SV12 = "SV12", SV21 = "SV21", SV28 = "SV28 complex", SV8 = "SV8 complex"))
```

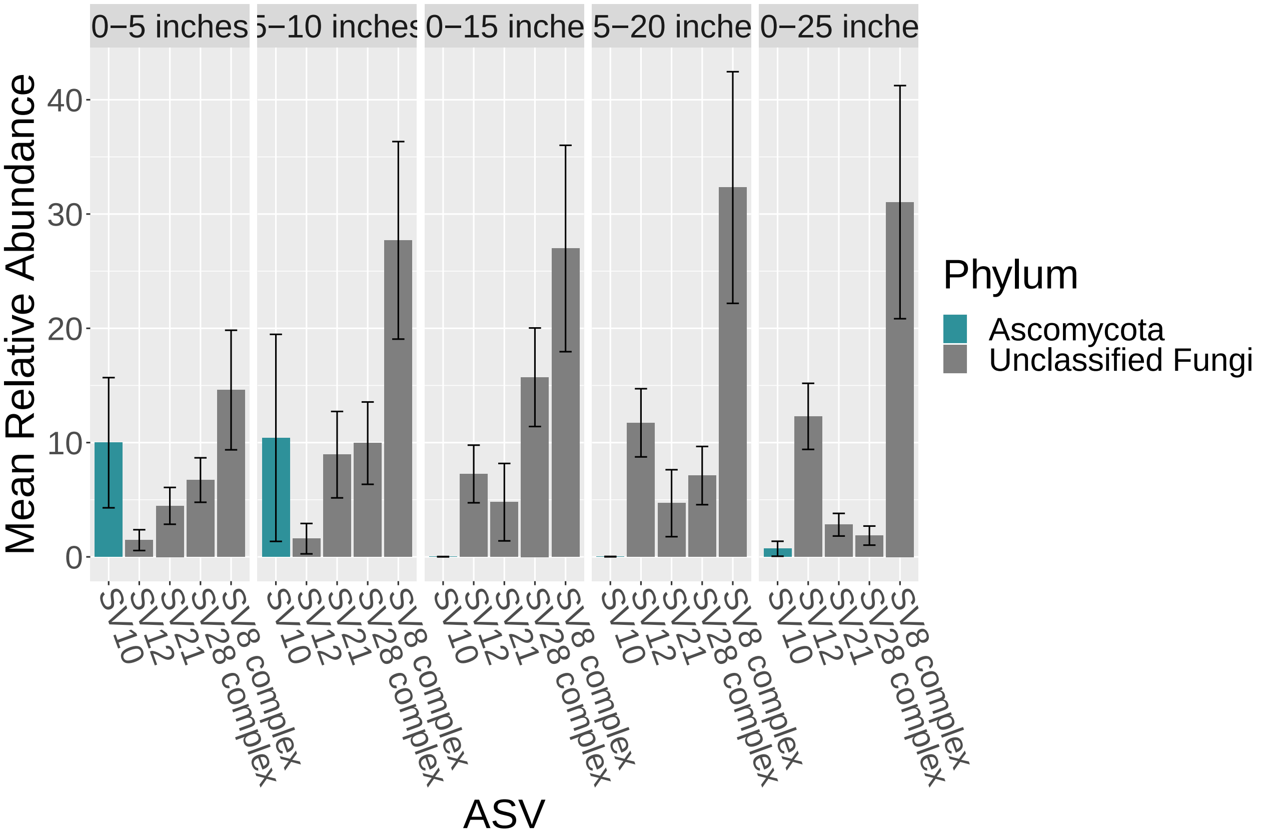


```
# write.csv(avgs_leaf,
# 'ps.nocontrols.pear_nomin_err3.COR.RA.NoNEG.rare5000.LEAF_ASVcomplex_mean02.csv'
# )
```

##### Stats on ASVs

```
# start
chisq = NULL
pvals = NULL
listofcats_sig = NULL
DataSet = df_leaf_asv

taxa = unique(DataSet$OTU)

for (cat in taxa) {
    new_df <- subset(DataSet, DataSet$OTU == cat)
    new_df$ST <- as.factor(new_df$SampleSpecificType)
    kw = kruskal.test(Abundance ~ ST, data = new_df)
    chisq = c(kw$statistic, chisq)
    pvals = c(kw$p.value, pvals)

    if (kw$p.value <= 0.05) {
        listofcats_sig = c(cat, listofcats_sig)
```

```
    }
}

pvals.bonf = p.adjust(pvals, method = "bonferroni")
df_taxa = data.frame(rev(taxa), chisq, pvals, pvals.bonf)
write.table(df_taxa, "LeafLength_rare5000_Mean_ASV_KW_02mean.txt",
    sep = "\t")


pvals.dunn = NULL
pvals.dunn.bonf = NULL
Zsc.dunn = NULL
comparison.dunn = NULL
cats.dunn = NULL

for (cat in listofcats_sig) {
    new_df <- subset(DataSet, DataSet$OTU == cat)
    new_df$ST <- as.factor(new_df$SampleSpecificType)
    dT = dunnTest(Abundance ~ ST, data = new_df, method = "bonferroni")

    for (i in 1:length(dT$res$Comparison)) {
        print(cat)

        pvals.dunn = c(dT$res$P.unadj[i], pvals.dunn)
        pvals.dunn.bonf = c(dT$res$P.adj[i], pvals.dunn.bonf)
        Zsc.dunn = c(dT$res$Z[i], Zsc.dunn)
        comparison.dunn = c(as.character(dT$res$Comparison)[i],
            comparison.dunn)
        cats.dunn = c(cat, cats.dunn)

    }

}


df_taxa = data.frame(cats.dunn, comparison.dunn, Zsc.dunn, pvals.dunn,
    pvals.dunn.bonf)
write.table(df_taxa, "LeafLength_rare5000_Mean_ASV_KW_02mean_Dunn.txt",
    sep = "\t")
```

##### Now split by epiphyte vs. endophyte status

```
# ASV by mean
AvgRA99_leaf = filter_taxa(ps.nocontrols.pear_nomin_err3.COR.rare5000.NoNEG.LEAF.RA,
    function(x) mean(x) > 0.02, TRUE)

# Lobulomycetales - SV8, SV11, SV16, SV56 Glomerellales -
# SV10 Aphelidiomycota - SV12 Chytridiomycetes - SV21
# Chytridiomycota - SV28, SV36

# SV8 complex = SV8, SV11, SV16, SV56 SV28 complex = SV28,
```

```r
# SV36

avgra99_leaf_merge <- merge_taxa(AvgRA99_leaf, c("SV8", "SV11",
    "SV16", "SV56"), "SV8")
avgra99_leaf_merge <- merge_taxa(avgra99_leaf_merge, c("SV28",
    "SV36"), "SV28")

df_leaf_asv2 <- psmelt(avgra99_leaf_merge)

grouped_leaf <- group_by(df_leaf_asv2, SampleType, SampleSubType,
    SampleSpecificType, OTU, Order, Class, Phylum)
avgs_leaf <- summarise(grouped_leaf, mean = 100 * mean(Abundance),
    sd = 100 * sd(Abundance), se = 100 * se(Abundance))

avgs_leaf %<>% mutate(Phyla2 = fct_explicit_na(Phylum, na_level = "Unclassified Fungi"))
avgs_leaf %<>% mutate(Order2 = fct_explicit_na(Order, na_level = "Unclassified Fungi"))
facet_names <- c(`0_to_5_inches` = "0-5 inches", `10_to_15_inches` = "10-15 inches",
    `15_to_20_inches` = "15-20 inches", `20_to_25_inches` = "20-25 inches",
    `5_to_10_inches` = "5-10 inches", Epiphyte = "Epiphyte",
    Endophyte = "Endophyte")
avgs_leaf$SampleSpecificType <- factor(avgs_leaf$SampleSpecificType,
    levels = c("0_to_5_inches", "5_to_10_inches", "10_to_15_inches",
        "15_to_20_inches", "20_to_25_inches"))

p = ggplot(avgs_leaf, aes(x = OTU, y = (mean), fill = Phyla2)) +
    geom_bar(stat = "identity", position = position_dodge()) +
    geom_errorbar(aes(ymin = (mean - se), ymax = (mean + se)),
        width = 0.4, position = position_dodge(0.9))
p = p + facet_grid(SampleSubType ~ SampleSpecificType, labeller = as_labeller(facet_names)) +
    theme(axis.text.x = element_text(angle = -70, hjust = 0,
        vjust = 0.5)) + ylab("Mean Relative Abundance") + theme(text = element_text(size = 28))
p + xlab("ASV") + guides(fill = guide_legend(title = "Phylum")) +
    scale_fill_manual(values = c("#2e919a", "grey50"), labels = c("Ascomycota",
        "Unclassified Fungi")) + scale_x_discrete(labels = c(SV10 = "SV10",
    SV12 = "SV12", SV21 = "SV21", SV28 = "SV28 complex", SV8 = "SV8 complex"))
```

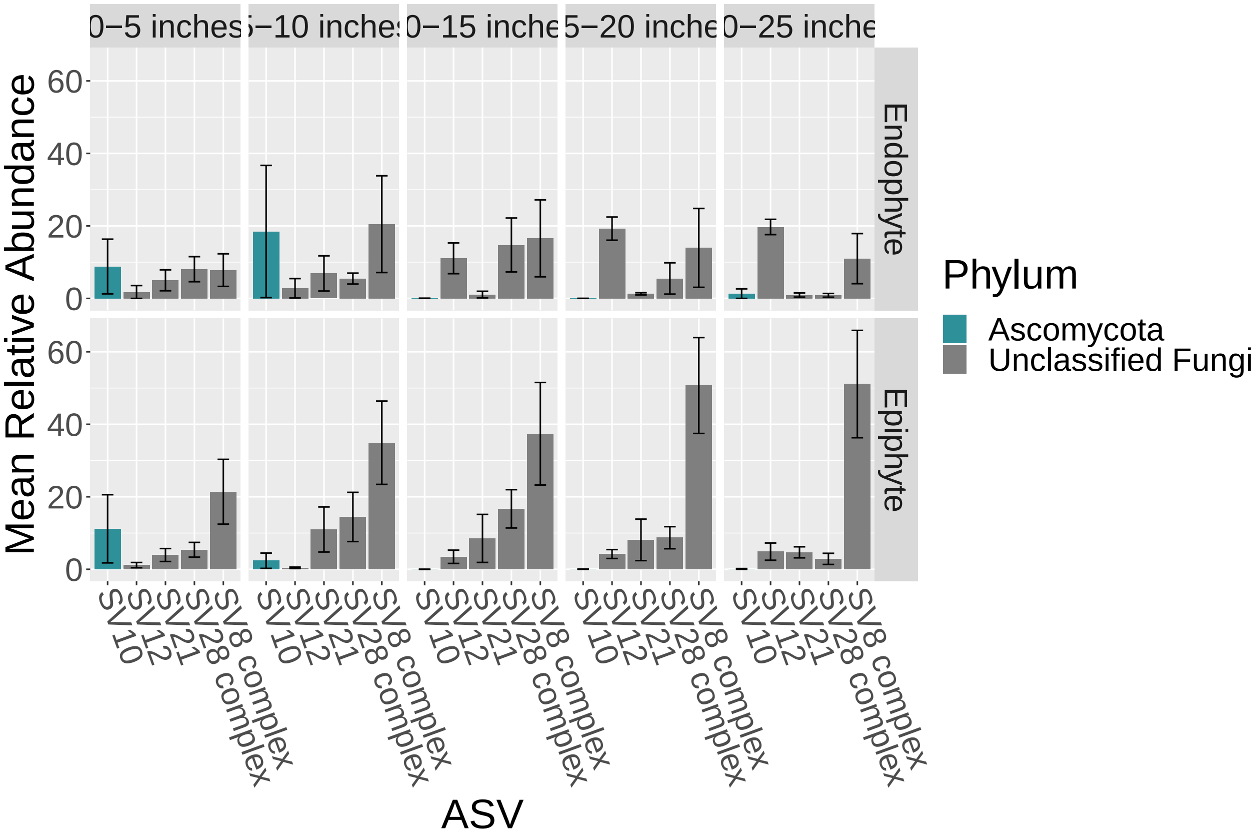


##### Stats on ASVs - are there ASVs that differ between epiphyte and endophyte leaf samples?

```
# start
chisq = NULL
pvals = NULL
listofcats_sig = NULL
DataSet = df_leaf_asv2

taxa = unique(DataSet$OTU)

for (cat in taxa) {
    new_df <- subset(DataSet, DataSet$OTU == cat)
    new_df$ST <- as.factor(new_df$SampleSubType)
    kw = kruskal.test(Abundance ~ ST, data = new_df)
    chisq = c(kw$statistic, chisq)
    pvals = c(kw$p.value, pvals)

    if (kw$p.value <= 0.05) {
        listofcats_sig = c(cat, listofcats_sig)

    }
}

pvals.bonf = p.adjust(pvals, method = "bonferroni")
```

```
df_taxa = data.frame(rev(taxa), chisq, pvals, pvals.bonf)
write.table(df_taxa, "LeafLength_EpiEndo_rare5000_Mean_ASV_KW_02mean.txt",
    sep = "\t")
```

##### Alpha diversity for leaf and root epi vs. endophytes

```
# leaf epi vs endo alpha
ps.nocontrols.pear_nomin_err3.COR.rare5000.NoNEG.NoRhizome <- subset_samples(ps.nocontrols.pear_nomin_e
    SampleType != "Rhizome")
ps.nocontrols.pear_nomin_err3.COR.rare5000.NoNEG.NoRhizome.NoSed <- subset_samples(ps.nocontrols.pear_n

    SampleType != "Sediment")

p = plot_richness(ps.nocontrols.pear_nomin_err3.COR.rare5000.NoNEG.NoRhizome.NoSed,
    measures = c("Observed", "Shannon"), x = "SampleType", color = "SampleSubType")
p = p + theme(text = element_text(size = 24)) + geom_boxplot() +
    geom_jitter() + theme(axis.text.x = element_text(angle = -70,
    hjust = 0, vjust = 0.5))
p + scale_colour_manual(values = c("#2e919a", "#fdc34a", "#3F5CAA",
    "grey50")) + xlab("") + guides(color = guide_legend(title = "Sample Type"))
```

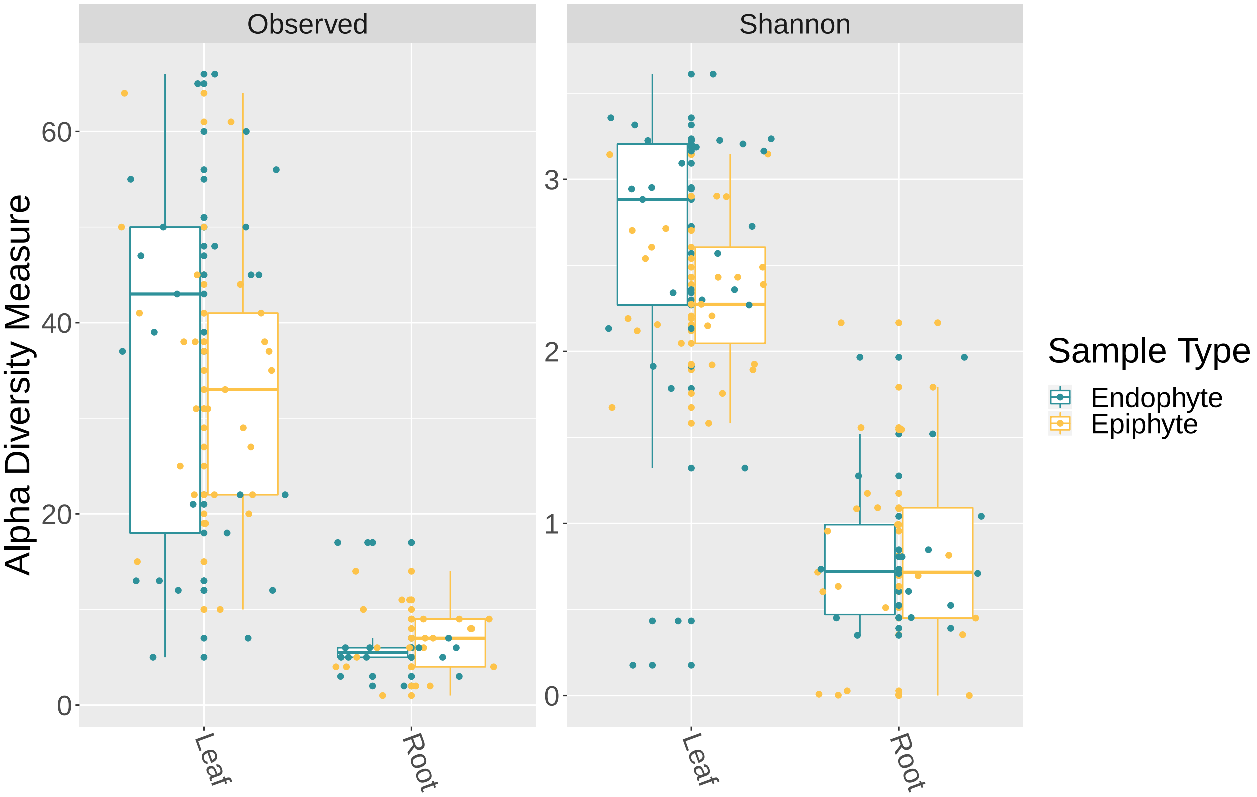


##### Stats on the alpha diversity

```
# Leaf epiphytes vs. endophytes

EE_Alpha = estimate_richness(ps.nocontrols.pear_nomin_err3.COR.rare5000.NoNEG.LEAF,
```

```r
    measures = c("Observed", "Shannon"))

EE_Alpha_2 <- cbind(EE_Alpha, sample_data(ps.nocontrols.pear_nomin_err3.COR.rare5000.NoNEG.LEAF))

EE_Alpha_2$ST <- as.factor(EE_Alpha_2$SampleSubType)

kruskal_test(Observed ~ ST, distribution = approximate(nresample = 9999),
    data = EE_Alpha_2)
```

```
##
##  Approximative Kruskal-Wallis Test
##
## data:  Observed by ST (Endophyte, Epiphyte)
## chi-squared = 0.462, p-value = 0.5112
```

```r
kruskal_test(Shannon ~ ST, distribution = approximate(nresample = 9999),
    data = EE_Alpha_2)
```

```
##
##  Approximative Kruskal-Wallis Test
##
## data:  Shannon by ST (Endophyte, Epiphyte)
## chi-squared = 4.3506, p-value = 0.0354
```

```r
# Root epiphytes vs endophytes

ps.nocontrols.pear_nomin_err3.COR.rare5000.NoNEG.ROOT <- subset_samples(ps.nocontrols.pear_nomin_err3.C
    SampleType != "Leaf")

EE_AlphaR = estimate_richness(ps.nocontrols.pear_nomin_err3.COR.rare5000.NoNEG.ROOT,
    measures = c("Observed", "Shannon"))

EE_Alpha_2R <- cbind(EE_AlphaR, sample_data(ps.nocontrols.pear_nomin_err3.COR.rare5000.NoNEG.ROOT))

EE_Alpha_2R$ST <- as.factor(EE_Alpha_2R$SampleSubType)

kruskal_test(Observed ~ ST, distribution = approximate(nresample = 9999),
    data = EE_Alpha_2R)
```

```
##
##  Approximative Kruskal-Wallis Test
##
## data:  Observed by ST (Endophyte, Epiphyte)
## chi-squared = 0.77689, p-value = 0.3914
```

```r
kruskal_test(Shannon ~ ST, distribution = approximate(nresample = 9999),
    data = EE_Alpha_2R)
```

```
##
##  Approximative Kruskal-Wallis Test
##
## data:  Shannon by ST (Endophyte, Epiphyte)
## chi-squared = 0.0011338, p-value = 0.9869
```
